# Not All Saliva Samples Are Equal: The Role of Cellular Heterogeneity in DNA methylation and Epigenetic Age Analyses with Biological and Psychosocial Factors

**DOI:** 10.1101/2024.11.08.621377

**Authors:** Meingold H. Chan, Mandy Meijer, Sarah M. Merrill, Maggie P. Y. Fu, David Lin, Julia L. MacIsaac, Jenna L. Riis, Douglas A. Granger, Elizabeth A. Thomas, Michael S. Kobor

## Abstract

Saliva is widely used in biomedical population research, including epigenetic analyses to investigate gene-environment interplay and identify biomarkers. Its minimally invasive collection procedure makes it ideal for studies in pediatric populations. Saliva is a heterogenous tissue composed of immune and buccal epithelial cells (BEC). Amongst the many epigenetic marks, DNA methylation (DNAm) is the most studied in human populations. Given DNAm’s integral role to cellular differentiation and maintenance, DNAm profiles are often highly cell type (CT)-specific and CT composition can drive salivary DNAm associations with environments or health as well as epigenetic age acceleration (EAA), discrepancy between chronological age and biological age derived from DNAm. To address this, reference-based CT deconvolution and statistically adjustment with estimated CT in DNAm analyses have become a common practice. However, it remains unclear how different CT reference panels—constructed from adult versus pediatric samples—affect DNAm results. Additionally, whether DNAm and EAA associations in saliva primarily originate from immune cells or BECs, or if they persist across saliva samples despite varying CT proportions, has yet to be examined.

The current study used salivary DNAm samples obtained from 529 children (mean age=7.26 years, SD=0.26 years) in a community-based cohort, the Family Life Project. Our results highlighted the impact of estimated CT discrepancies across child and adult reference panels on DNAm associations. Upon stratifying the salivary DNAm samples into three subsamples—primarily BECs, primarily immune cells, and an approximately equal mix of both, we found significantly different EAAs across stratified samples when CT proportions were not accounted for. In both the context of DNAm and EAA associations, we detected stronger effects of cotinine concentrations, a tobacco smoke-exposure biomarker, in the subsample with primarily immune cells. We discussed the implications of our findings for the interpretation and replication of epigenetic research involving pediatric saliva samples.

---

Epigenetics has gained increasing interest as a potential biological mechanism through which environmental and psychosocial factors can influence health and development. DNA methylation (DNAm) is a key epigenetic modification that is well-characterized and widely studied in humans for its potential to both act as a biomarker and elucidate pathways of risk and resilience^1^. DNAm studies are typically conducted using epigenome-wide association studies (EWAS) to identify DNAm links to phenotypic and environmental differences. More recently, epigenetic clocks have emerged as a major tool to harness information of the methylome, comparing DNAm-derived biological age of cells to the chronological age of the individual.

Irrespective of the epigenetic studies’ approaches, these DNAm association analyses need to contend with the fundamental role of DNAm in cellular differentiation and maintenance. The latter often result in cell type (CT)’s having distinct DNAm profiles^2^. Given that the majority of human DNAm research involves peripheral bulk tissues, which are often composed of multiple different CTs, it is an important consideration to account for cellular heterogeneity in statistical analyses. To address this issue, CT proportions can be quantified by cell count or bioinformatically estimated from CT reference panels of sorted cells’ DNAm^3,4^.

Oral samples, including saliva, are often preferred in biosocial and epidemiological research involving children as the collection methods are minimally invasive and more readily accepted by parents^5,6^. Additionally, samples can be collected at home and mailed at room temperature, resulting in lower technical demands^5^. These unique strengths of saliva samples have generated considerable opportunities in pediatric DNAm investigations, yet its high CT heterogeneity presents an analytical challenge that needs to be addressed to ensure rigorous DNAm analyses and downstream interpretation^2^.

Saliva is composed of both buccal epithelial cells (BECs) and immune cells^5^. Its cellular composition is highly variable across individuals^3,7,8^, which can be attributed to wide ranging factors, including social exposures, oral hygiene, salivary collection methods, and, most importantly, age. Therefore, estimating CT proportions using reference panels trained on samples of different tissues and developmental stages (i.e., pediatric vs. adult) could generate differential outputs. These discrepancies in CT proportion estimation could also impact downstream DNAm analyses and the identification of differentially methylated sites in saliva^2,3^.

Furthermore, deeper investigation of how the heterogeneity in saliva may influence the interpretation and reproducibility of DNAm results is needed. Considering saliva samples with extreme interindividual variation of BEC proportions, one could reason that DNAm associations identified in saliva samples with extremely *low* BEC proportion (and therefore extremely *high* immune cell proportion) will be more similar to that in tissues of primarily immune cell composition, like blood, though these immune cells are oral in origin. In contrast, those with extremely *high* BEC proportion may have DNAm associations more similar to tissues with a largely epithelial composition, such as cheek swabs. Therefore, DNAm associations that can be identified with salivary samples may be highly dependent on the CT compositions in the specific samples and these differential associations may parallel tissue differences. This leads to challenges in the interpretation and replication of DNAm associations found in saliva samples that are considered the same tissue source yet may have highly distinct CT proportions.

Some variables of interest may be more likely to have differential DNAm associations across salivary samples of differing primary CT populations, with specific bias towards a particular CT. For example, exposure to tobacco smoke may exhibit stronger associations with DNAm in immune cells than BECs in saliva since it is a major source of oxidative stress and could influence the mucosal immune system and in turn affect systemic immune responses^9^. Thus, samples with higher abundance of immune cells may have higher sensitivity to smoke exposure. This high CT heterogeneity in salivary samples complicates interpretation, replication, and meta-analyses of DNAm results. Nevertheless, such heterogeneity may also serve as an opportunity to elucidate what CTs may underpin salivary EWAS or EAA associations, if any in particular.

Relatedly, salivary CT proportions may also influence the biological age estimated by epigenetic clocks trained in specific tissues and populations using DNAm, known as epigenetic age. Indeed, epigenetic age acceleration (EAA) derived from the deviation between epigenetic and chronological age has been shown to be affected by CT proportions in cheek swab samples^10^. Given the developmental changes in BEC proportion in pediatric saliva and the previous findings on the association between EAA and BEC proportion^10^, it is reasonable to expect that saliva samples with differential BEC proportions may show faster or slower epigenetic aging. Thus, it may be pertinent to account for CT proportions when calculating EAA to reduce confounding effects.

In the current study, we leveraged salivary samples from 529 7-year-old children in a community cohort with a naturally-occurring highly variable CT proportions, similar to that previously shown in other published samples (i.e., 0-100% estimated BEC^3^ and 20-100% from count data^7^). Given that reference panels may affect estimation of CT proportion and thus DNAm analyses, we characterized and compared CT proportions predicted with the adult and child reference panels and explore its effect on EWAS outcomes. While saliva has been traditionally analyzed together as a whole, we characterized the specific CT populations underpinnings of DNAm associations (both EWAS and EAA) in heterogeneous salivary samples by stratifying these samples based on primary CT. This could help elucidate if DNAm signals observed in saliva originated primarily from immune cells or BECs or can be found across saliva samples despite differing CT proportions. To maximize the applicability of our findings across disciplines, and to illuminate the importance of choosing an appropriate reference for DNAm analysis, we chose three variables that are commonly of interest in biosocial research, spanning across the spectrum of biological to social. These included biological sex, a variable of primarily biological influence; salivary cotinine concentration, which is indicative of tobacco smoke exposure and represents a combination of biological and social influences, and socioeconomic status (SES), a variable of primarily social influence. These variables have been widely studied in the context of DNAm across disciplines^9,11–13^. They also represent varying degrees of expected CT-specific DNAm associations.

## Methods

### Participants

The current study used archival saliva samples collected as part of the Family Life Project’s (FLP) 7-year follow-up assessment. As previously described^14^, a representative sample of 1,292 families were originally recruited at the time that mothers gave birth. A subset of 743 FLP families provided consent for analysis of the saliva samples at the 7-year follow-up. After quality control (QC) procedures, DNAm data from 529 children (mean age=7.26 years, SD=0.26 years, 49.9% boys, mother-reported child race: 59.5% White and 39.7% Black/African American) and survey data from their primary caregivers (98.4% biological parents) were used in our analyses (**Supplementary Table S1**).

### Parent-reported measures: Children’s sex & socioeconomic status

Primary caregivers reported on children’s biological sex at birth, which was confirmed by the DNAm data in the final samples used for analyses (described in *DNAm preprocessing* below). Income-to-needs ratio, as an index of SES, was calculated by dividing parent-reported total household income by the federal poverty threshold for the year 2011, adjusted for the number of persons in the home. An income-to-needs ratio of 1.00 or below indicates family income at or below the poverty level, adjusted for household size^14^.

### Salivary cotinine measure

Salivary cotinine was assayed on whole saliva samples collected from children in duplicate using a commercially-available, ELISA kit (Salimetrics LLC, Cat ##1-2002, Carlsbad, CA) following the manufacturer’s protocol and per our previous studies^15^ (See **Supplementary Material** for details). Concentrations below the lower limit of detection (LLD; n=135) were treated as missing data in the main analyses.

### Salivary DNAm profiling and preprocessing & Epigenetic Age calculation

At the same visit as salivary cotinine collection measurement, whole saliva samples, from which DNA was extracted, were also collected from children (see **Supplementary Material** for details on collection of saliva sample and DNA extractions). DNAm profiling by the llumina MethylationEPIC BeadChips (EPIC array, Illumina, CA, USA) was performed at the University of British Columbia (Vancouver, BC, Canada) as previously described^16^. Briefly, QC of the EPIC array DNAm data was performed in R (4.2.2) using a combination of the minfi package (v1.44.0)^17^ and ewastools (v1.7.2)^18^. A total of 200 samples were removed after all QC steps (**Supplementary Figure S1a**). Four additional samples with age mismatch (>10 years difference) between chronological and epigenetic age were removed. based on the epigenetic age calculated with the Pediatric Buccal (PedBE)^16^ and Horvath Skin-Blood clock^19^ in the methylclock R package and reported chronological age were removed. A total of 529 high-quality samples were included for downstream analyses (**Supplementary figure S1a**). Probe filtering was conducted on the normalized DNAm data. A total of 786,069 probes passed the QC.

### Whole genome genotyping & genetic principal components

Illumina Global Screening v1.0 arrays (Illumina, CA, USA) were used to measure 654,027 single nucleotide polymorphisms (SNPs) for every individual. Genetic ancestry principal components (PCs) were computed from SNPs^20^. To adjust for effects of genetic admixture, we included the first three PCs to in all analyses.

## Statistical analyses

### Characterizing and comparing CT proportion estimates across adult and pediatric saliva reference panels & its effect as covariates on EWAS outcomes

First, we estimated the CT proportions of BECs and immune cells with robust partial correlation with both an adult^21^ and a pediatric saliva-based reference panel^3^. The first two PCs of our CT proportion variables explained>90% of the variance (adult: 94.6% and child: 97.2%) and were included in the EWAS models.

A total of 614,686 variable probes, where the DNAm level (β value) varies by at least 5% across samples in the 5th and 95th percentile, were included for the EWASs. Two EWAS models were run for each variable of interest, adjusting for different estimated CT proportions, either with the pediatric or adult CT reference panel (six EWAS models in total). The other covariates for both models were the same—the first three genetic PCs and child’s age. The robust linear regression model used for the EWAS was:

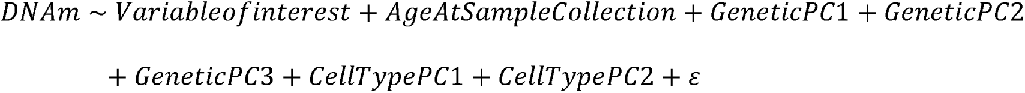

Differences in DNAm associated with the variable of interest was quantified by the difference in beta value (Δβ)^1^. We used the Benjamini-Hochberg False Discovery Rate (FDR) control method for multiple-test correction^22^. A statistical threshold was set at FDR<0.05 to determine significant associations between DNAm at each site with each variable of interest. We also set a technical threshold of |Δβ|>.035, which was greater than technical noise from the DNAm array. Across the two reference panels, we compared the sites that were identified to be statistically significantly associated with each variable of interest.

### Comparing EWAS results between stratified samples by BEC proportions

Using a data-driven approach, we stratified the samples into three groups–low-, mid-, and high BEC groups, consistently suggested by both Jenk’s natural break^23^ and k-mean clusters^24^. With the same three variables of interest, one EWAS for each variable was run with each stratified subsample (i.e., nine EWAS models). We reported the unique number of significant DNAm associations from each subsample and the number of overlapping sites across the three subsamples. Since the sample sizes of the three subsamples were unequal, we ran 100 randomized permutations in which we randomly assigned individuals to one of the three BEC groups, independent of estimated BEC proportions. The resulting permutation p-value represents the likelihood that a smaller number of associations are observed with a smaller sample size (see **Supplementary Material** for details and results).

Sensitivity analyses were conducted for sex associations to examine cross-tissue comparison using an independent cohort as well as for cotinine to address the potential impact of missing data and categorizing the cotinine concentration into a three-level variable on the results and (see **Supplementary Material**).

### Comparing EAAs and their associations across stratified samples by BEC proportions

We compared PedBE and Horvath Skin-blood EAA calculated with and without adjusting estimated CT proportions across the three stratified samples using ANOVA. Additionally, we examined the associations between EAA (with and without CT adjustments) with the same three variables of interest in the full sample and the three stratified samples. We used ANOVA for biological sex, robust linear regression for cotinine concentration given its skewness, and simple linear regression for SES.

## Results

### The child reference panel was more appropriate than adult panel for pediatric salivary CT proportion estimation

Using a large cohort of children’s saliva, our study addressed several fundamental questions relevant to pediatric salivary DNAm analyses (**Figure 1**). Our initial focus was to test the performance of existing adult and child reference panels to estimate CT proportions in saliva. With both reference panels, BEC proportions were estimated to be the highest and the most variable among all CT proportions (**Figure 2a; Supplementary Figure S2**). The estimated proportion of immune cells, primarily neutrophils, had a significant and strong negative correlation with estimated BEC proportion, *r*(527)=-0.93, p<2.2e-16). Therefore, estimated BEC proportion was likely to capture most of the variability of estimated CT proportions in saliva and were used in our remaining analyses.

**Figure 1.**
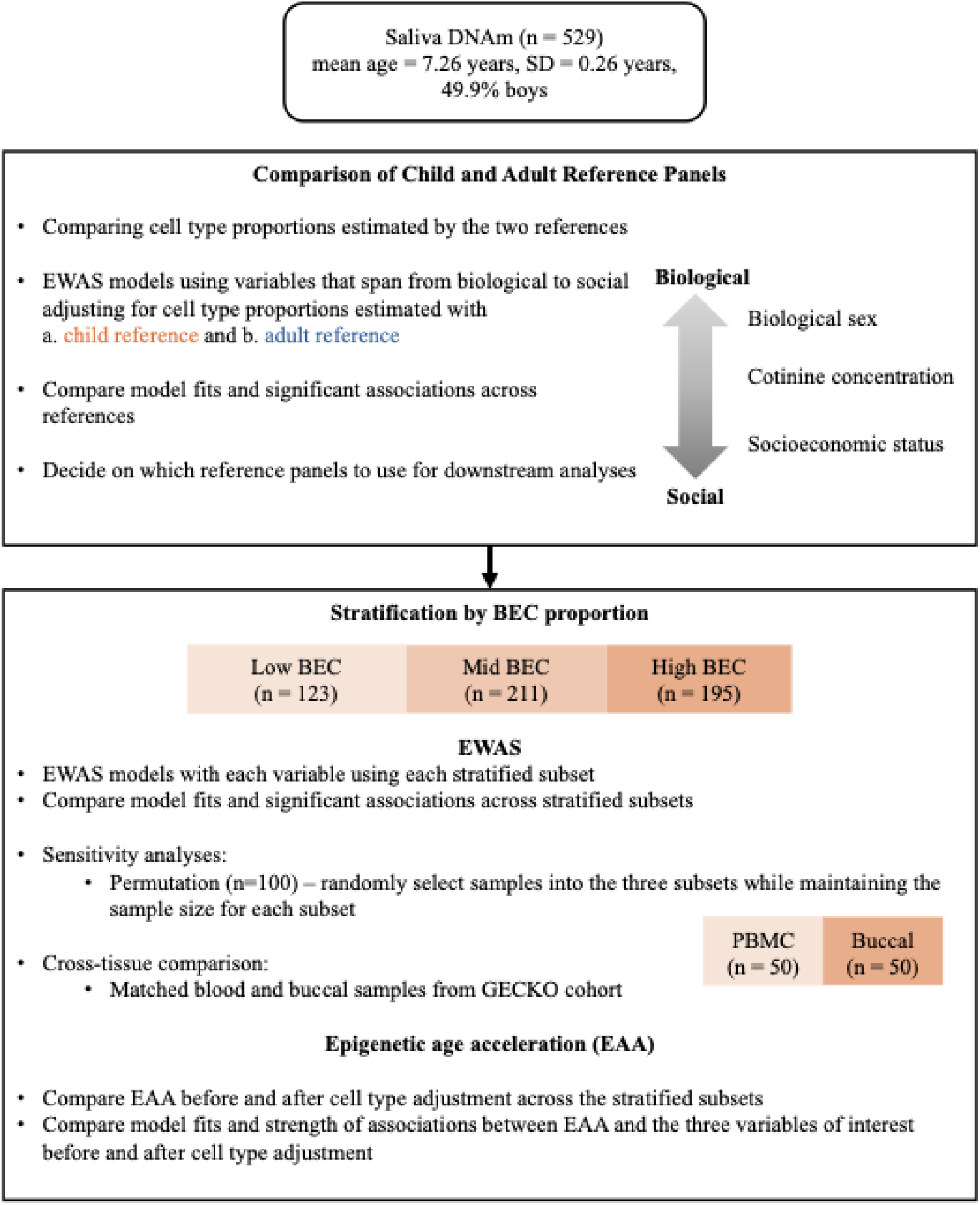
Study Overview. Cell type (CT) proportions were estimated with both a child and an adult salivary reference panel. The first aim of the study was to compare and characterize the estimated CT proportions across reference panels and examine the effect of CT proportions estimated from different reference panels on EWAS results of biological sex, cotinine concentration, and socioeconomic status. The second aim was to investigate the effect of stratification by buccal epithelial cell proportion on EWAS and EAA associations. CETYGO = CEll TYpe deconvolution GOodness.

**Figure 2.**
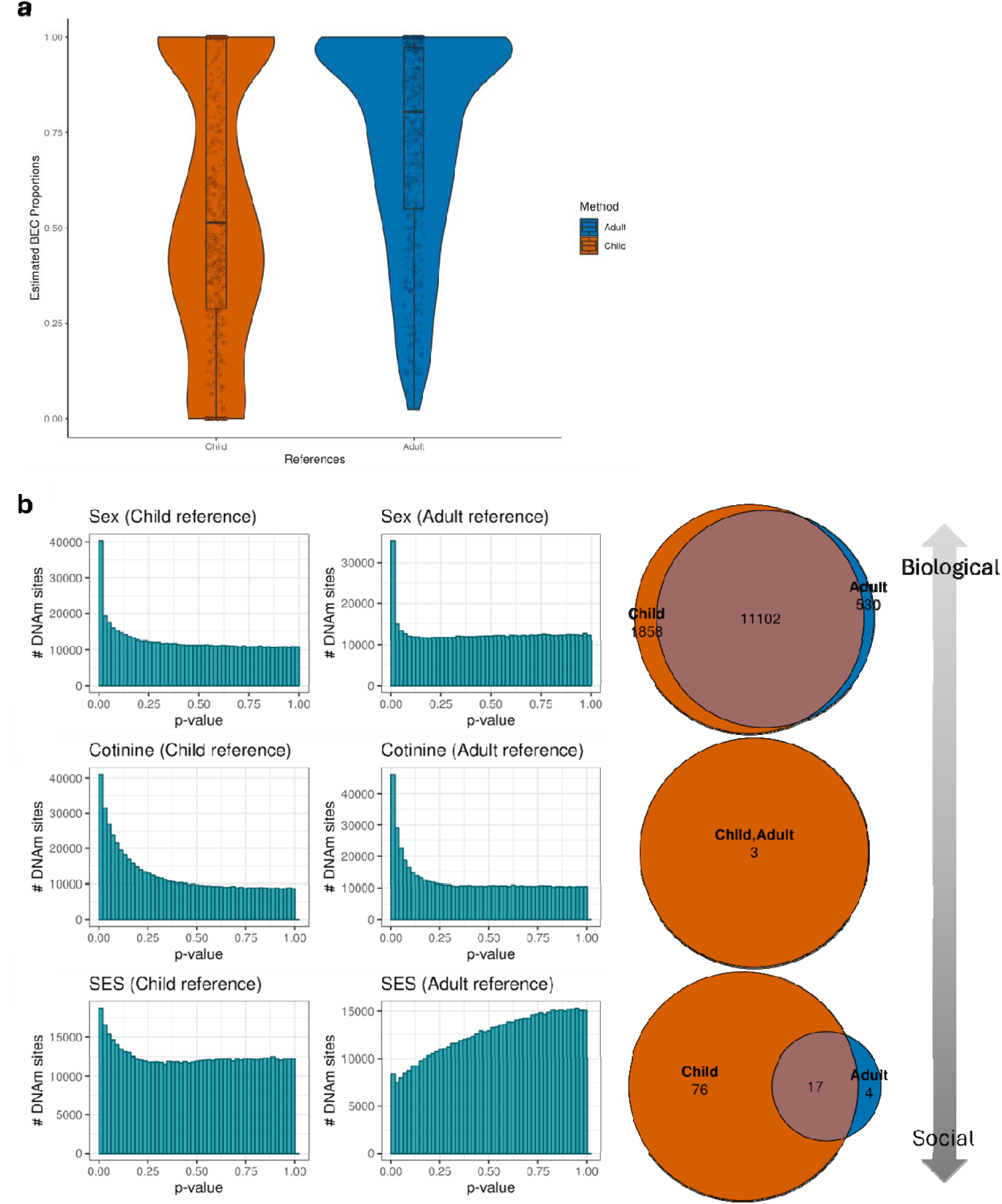
Estimated buccal epithelial cell (BEC) proportion and significantly associated DNAm sites differed across child and adult reference panels. (a) A violin plot of BEC proportions in pediatric saliva as estimated with child and adult reference panels using the DNAm-based deconvolution tool EpiDISH. Orange (left) reflects the child reference panel, and blue (right) reflects the adult reference panel. Estimated BEC proportion means were different as indicated by paired-sample t-test, *t*(528)=31.788, *p*<2.2e-16. (b) Each row shows the results of each variable of interest (sex, cotinine concentration, and socioeconomic status) across the child and adult reference panels. This includes the p-value histograms of the EWASs and Venn diagrams representing the unique and overlapping sites associated with the variable of interest. The range of Δβ extracted from EWAS (with child-reference estimated CT proportions) varied across variables (sex: -0.394 to 0.290, cotinine: -0.095 to 0.090, SES: -0.0967 to 0.147).

The estimated BEC proportions were significantly correlated across reference panels (Pearson’s *r*=0.950, *p*<2.2e-16). The child reference panel estimated a lower median of BEC proportion (51.34%) and higher IQR (71.18%) than the adult reference panel (median: 80.46%; IQR: 42.29%). The CETYGO score, with lower score indicating higher appropriateness, of the child reference panel was below 0.1 (mean=0.074, range=0.03-0.09), while the CETYGO score of the adult reference panel was above 0.1 (mean=0.15, range=0.05–0.20), suggesting that the child reference panel is a more appropriate panel for pediatric samples than the adult reference panel. Therefore, all downstream analyses were conducted with the child panel.

### Reference panels produced different outcomes for an EWAS of SES and similar results for biological sex and cotinine

We then tested whether the observed discrepancies in estimated CT proportions from the two reference panels influenced EWAS results. First, we note that a larger number of DNAm associations were identified with biological sex than cotinine concentrations and SES with both references. Second, across references, EWAS for sex and cotinine concentration had high overlaps in statistically significantly associated sites (FDR<.05 and |Δβ|>.035, 92% and 100%, respectively) and comparable model fits for estimations (**Figure 2b; Supplementary figure S3**). Sensitivity analyses with imputed cotinine concentration and low vs. moderate exposure also showed strong overlaps in statistically significantly associated sites in child reference panel as compared to adult reference panel (85% and 100%, respectively) (**Supplementary figure S4**).

However, differences in DNAm associations with SES were observed across reference panels (only 18% overlap), with the model adjusted for the child reference estimated CT proportions having a higher number of unique significant DNAm associations than adult reference (76 and 4, respectively) and more well-behaved p-value histogram (**Figure 2b; Supplementary figures S3 and S4**).

### Stratification by estimated BEC proportion detected a larger number of significant DNAm associations with sex and cotinine concentrations, but not SES

Our findings of the distinct trimodal distribution of estimated BEC samples with child salivary reference suggested that stratification of the samples may provide us insight into the particular CT composition in which salivary DNAm signals can be detected, both in the context of EWAS and EAA. Thus, we stratified the samples into three groups and repeated the analyses on these subsamples, herein described as high, mid, and low BEC subsamples, which contained primarily BEC, primarily neutrophils, or an approximately even mix of both. We characterized the three subsamples by comparing their demographic characteristics using one-way Analysis of Variance (ANOVA) and confirmed that the three stratified subsamples were not different at the demographic level, including child sex, age, parent-reported race, SES, and cotinine concentration based on one-way ANOVA, *p*s=0.38-0.97.

EWAS for biological sex performed in the high and low subsamples, showed an increased number of significant associations (n associations=3,007 and 1,980, respectively), compared to the full sample (n associations=1,780) (**Figure 3; Supplementary Figure S5a**). However, the medium BEC subsamples, despite having the largest sample size, had the least number of sites associated with sex (n associations=1,422). We observed 11.9% overlapping significantly associated sites (559 sites) across the three stratified subsamples of different levels BEC proportions (**Figure 3**). To compare the extent to which these differential EWAS outcomes across stratified subsamples may be akin to tissue effect, we leveraged the matched buccal and PBMC samples from GECKO, our validation cohort. We showed a lower percentage of overlapping DNAm associations with sex across tissues (5%, **Supplementary Figure S6**) than that across the three BEC subsamples in our main analyses with the discovery cohort.

**Figure 3.**
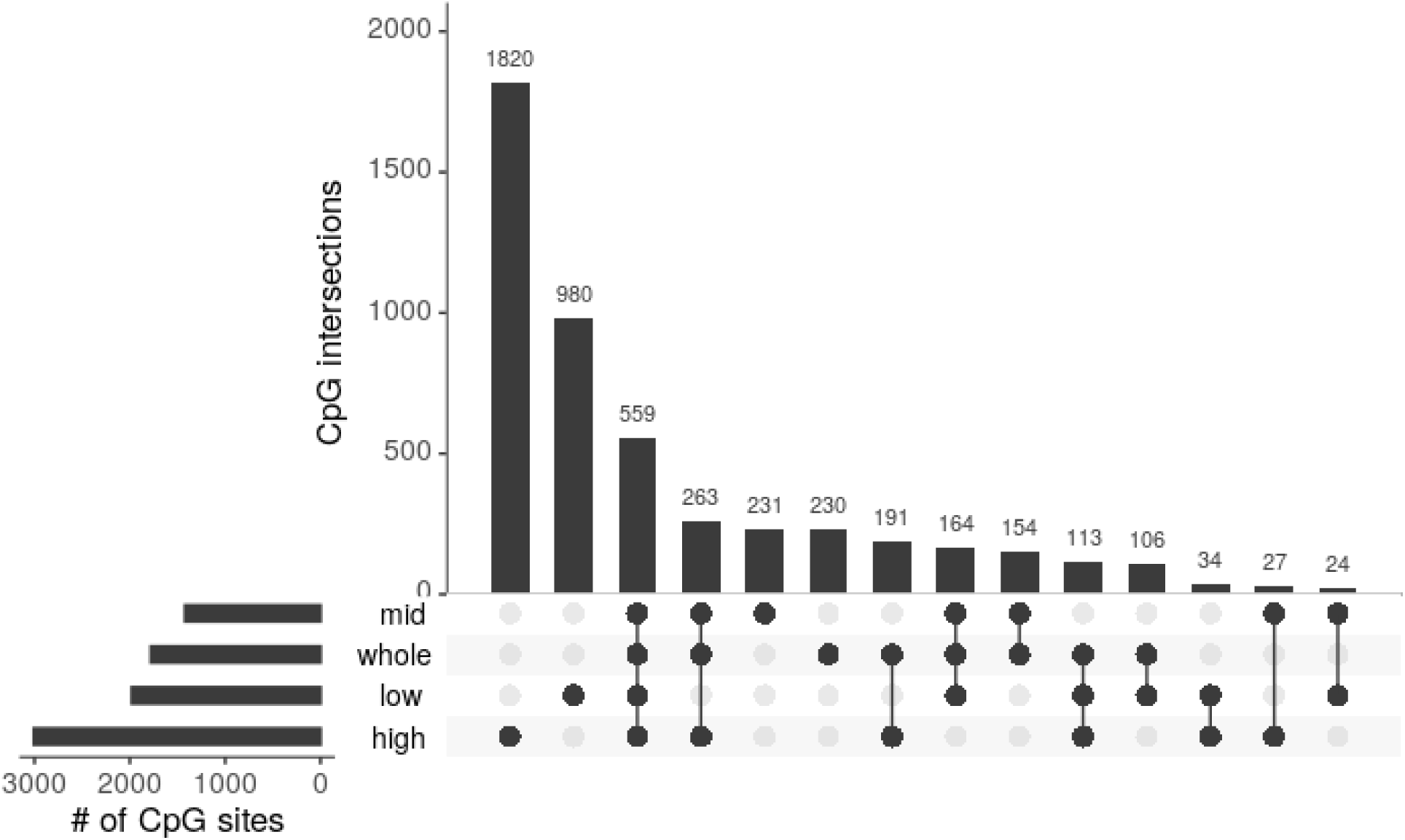
Significant associations with sex overlapping across stratified and full samples. EWAS for sex in the full sample as well as the high (n=195), mid (n=211), and low (n=123) BEC subsamples adjusting for CT proportions estimated by the child reference panel. The upset plot shows the unique and overlapping significant associations in the three stratified samples and the full sample.

We observed significant associations with cotinine concentration only in the low BEC group in both the main analysis (**Supplementary Figure S5b)** and the sensitivity analysis with imputed cotinine concentrations (**Supplementary Figure S7a**). Despite having a smaller sample size, a greater number of sites were significantly associated with cotinine concentrations in the low BEC subsample compared to the full sample (n associations=784 vs. 3, respectively).

Lastly, we did not find any significant associations with SES when the samples were stratified by BEC proportions (**Supplementary Figure S5c**). To assess whether the directionality of the effects in SES-associated sites identified with the full sample were consistent in the stratified samples, we conducted Pearson’s correlation test on Δβ across full and stratified samples (**Supplementary Figure S8**). While the overall effect sizes were smaller in the stratified samples than the full sample, the Δβs of these sites were strongly and positively correlated across full and stratified samples, *r*s=0.81-0.96, *p*s < 2.2e-16.

### Epigenetic Age Acceleration differed across BEC stratified samples without regressing out CT proportions

EAA, which reflects the discrepancy between EA and chronological age, can be affected by CT proportions^10^. To test for potential CT effect in saliva, we compared EAA calculated with PedBE and Horvath Skin-blood clocks across stratified samples, with and without CT adjustment in EAA calculations. We calculated the EAAs for both clocks by extracting residuals from the linear models epigenetic age regressed on 1) only chronological age and 2) both chronological age and estimated BEC proportions.

When CT proportions were not adjusted for in EAA calculations, both PedBE and Horvath Skin-blood EAAs were significantly different across BEC stratified subsamples, *F*(2, 526)=11.6, *p*=1.17e-05 and *F*(2, 526)=32.69, *p*= 4.16e-14, respectively (**Figures 4a & c)**. Specifically, children in the high BEC subsample showed lower EAA without CT adjustment than the other subsamples. These differences disappear when CT proportions were adjusted for in EAA calculations, *p*s=0.27-0.93 (**Figures 4b & d**).

**Figure 4.**
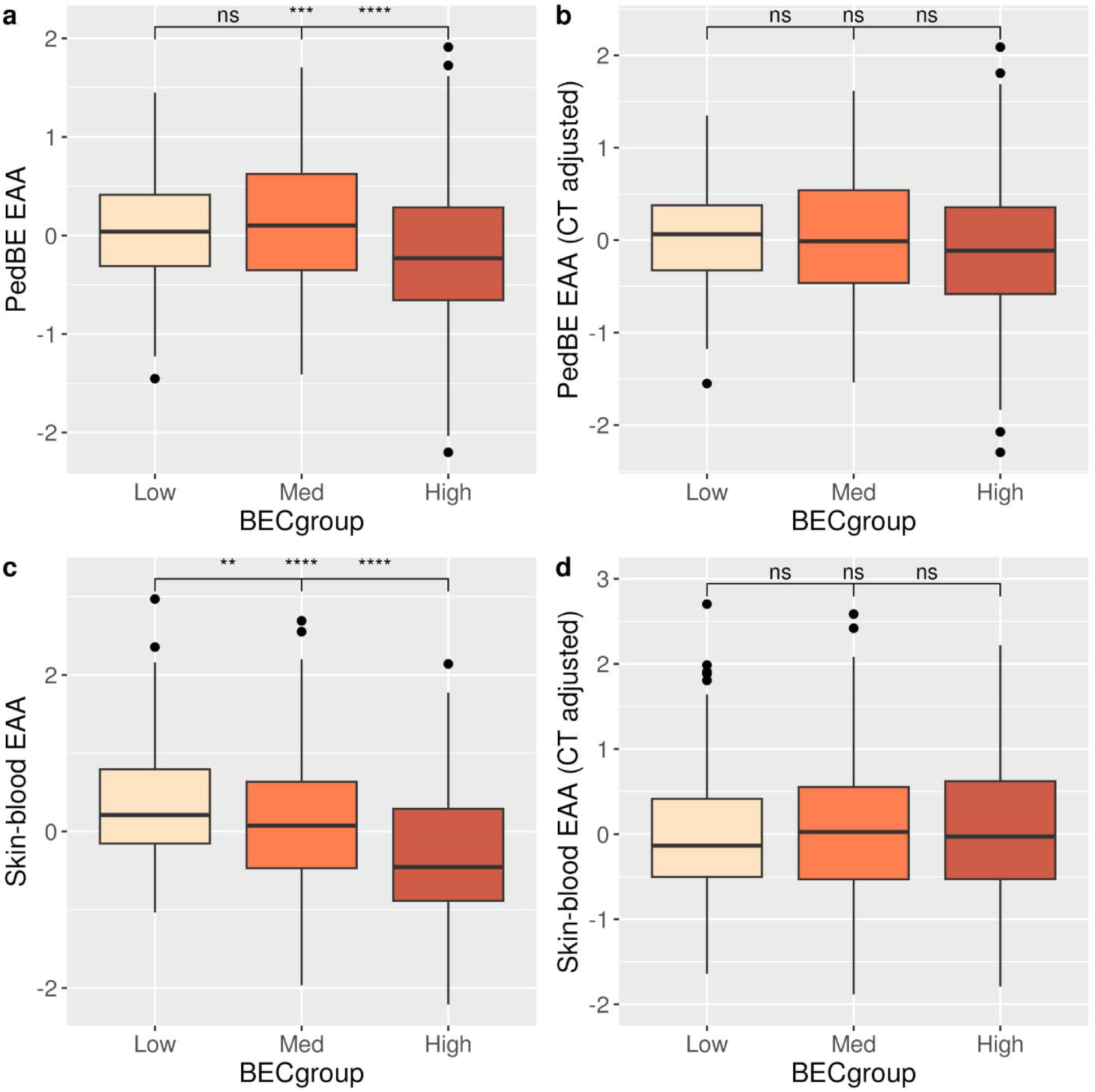
Significant differences in PedBE and Horvath Skin-blood EAA across BEC groups without CT adjustments but not with CT adjustment. Boxplot showing the PedBE EAA of the three BEC groups when calculated a) without CT adjustment and b) with CT adjustment and that of Horvath Skin-blood EAA c) without CT adjustment and d) with CT adjustment. Children in the high BEC subsample showed lower EAA without CT adjustment than the other subsamples (-0.31 and -0.24 years in PedBE EAA and -0.44 and -0.72 in Skin-blood EAA).

### Stratification by BEC proportions and adjusting for CT proportions in PedBE EAA calculations change the associations with variables of interest

Given the observed impact of CT proportions in EAA calculation and the common usage of EAA as a biomarker reflecting exposures and health outcomes in the field, we investigated whether PedBE EAA associations with our variables of interest were impacted by the estimated CT proportions in EAA calculation as well as stratification by estimated BEC proportions. The same set of analyses were also conducted with the Horvath Skin-blood EAAs with null associations in all conditions (**Supplementary Figure S9**).

Both PedBE EAAs with and without CT adjustment showed significant sex difference in the full sample, *F*(1, 527)=7.70, *p*=0.0057 and *F*(1, 527)=9.67, *p*=0.0020, respectively. Specifically, we found lower EAA in boys than girls with both EAAs with CT adjustment and without and similar effect size (-0.18 and -0.16 years, respectively) (**Figure 5a)**. After stratification, sex difference in both PedBE EAAs were observed only in the mid BEC subsample, *F*(1, 209)=6.33, *p*=0.0126 and *F*(1, 209)=6.336, *p*=0.0125, respectively (**Figure 5a)**. Again, boys showed lower EAA than girls, with similar effect size for EAA adjusted and not adjusted for CT proportions (-0.23 and -0.22 years). It is worth noting that the overall effect size in the mid BEC subsample was stronger than in the full sample.

**Figure 5.**
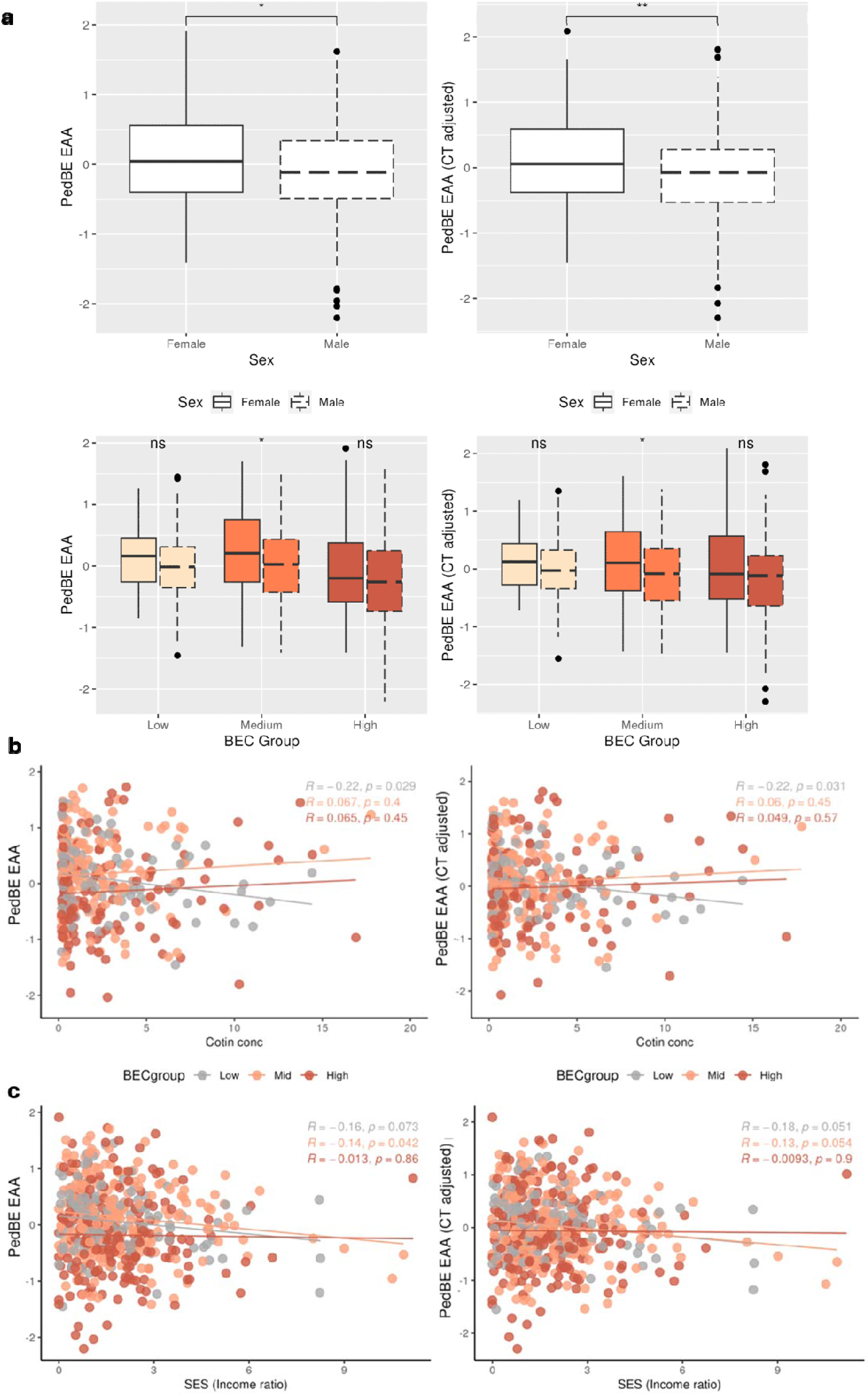
PedBE EAAs significantly differed across biological sex and associated with cotinine concentration in specific BEC stratified subsets. Associations between variables of interest and two PedBE EAAs—regressed onto chronological age only and regressed onto both chronological age and CT proportions—were shown. Panel *a* showed boxplots of PedBE EAAs across biological sex (male = 1, female = 0) in each stratified BEC subset, showing significantly lower EAA in boys than girls only in the mid BEC subset. Panel *b* and *c* showed scatterplots with regression lines (low BEC in grey, mid BEC in orange, and high BEC in brown) depicting the associations between PedBE EAAs and cotinine concentration (Cotin conc) and SES (income ratio), respectively. PedBE EAA were significantly associated with Cotin conc only in low BEC subset (both EAA variables) and SES only in mid BEC subset when CT proportions were not regressed out in EAA calculation.

While both PedBE EAAs were not significantly associated with cotinine concentrations in the full sample (*p*s=.391 and .456), after stratification we found a significant negative association in the low BEC subsamples between cotinine concentrations and both EAAs with and without CT adjustments, *r=*-0.22*, p=*0.029 and *r=*-0.022*, p=*0.031, respectively (**Figure 5b**).

Lastly, in the full sample, the association between SES and EAA without accounting for CT proportions was not statistically significant (*p*=0.054) but became statistically significant (p=0.039) using EAA accounted for CT proportions. Specifically, higher SES was associated with lower EAA, *r=*-0.09*, p=*0.039. Among stratified samples, the same association was found in the mid BEC group only with EAA not accounting for CT proportions, *r=*-0.14*, p=*0.042 (**Figure 5c**).

## Discussion

The current study addressed the challenges in analyzing and interpretating DNAm findings of high CT heterogeneity pediatric saliva samples both in the context of EWAS and EAA associations using salivary DNAm samples from a large cohort of children. Capitalizing on the high heterogeneity nature of saliva, we investigated whether there are CT-specific associations in BEC or immune cells in DNAm and EAA associations using a stratification approach while adjusting for estimated CT proportions calculated with the appropriate reference panel. Overall, our study demonstrated that a pediatric-specific saliva reference panel provided more appropriate CT proportion estimations than an adult reference and stratifying samples by BEC proportion enhanced sensitivity in detecting DNAm and EAA associations in multiple variables of interest. These findings provide critical guidance for improving the interpretability and rigor of epigenetic analyses using pediatric saliva samples.

### Reference panel choice influenced estimated BEC proportions, having differential impact on DNAm associations with biological and social variables

Our CT estimation results were highly comparable with findings from the previous study that generated the pediatric saliva reference panel^3^, showing lower and more variable estimated BEC proportions than the adult reference. The wider range of estimated BEC proportions with the child reference compared to the adult reference is likely reflective of natural biological variation of pediatric saliva as indicated by a highly variable CT proportion observed with actual cell count data^3,7^. Furthermore, our study demonstrated that discrepancies in CT estimation differentially impacted the outcomes of statistical models investigating high dimensional DNAm associations with variables that span across the biological and social spectrum. Specifically, we found that EWAS results of a social variable, SES, was the most impacted by the estimated CT discrepancies across references. Although the current analyses could not empirically differentiate true positive from false positive findings, it is tempting to speculate that adjusting for potentially more appropriate CT estimations generated in data of a similar tissue and age range was able to better capture variability of DNAm associated with CT and increased the power to detect significant associations with SES^25^. Yet, it is unclear why such impact was only observed with the most biologically distal and socially proximal variable in our study. Future investigations on other social variables are needed to scrutinize the effect of the CT estimation discrepancy. Nonetheless, our findings add to the growing body of literature creating awareness around the considerations of CT proportions in DNAm analyses^1,26^.

### Stratified samples by BEC proportion allowed detection of tissue-specific effects in EWAS

Interpreting and replicating DNAm findings from an extremely heterogenous tissue like saliva is challenging as it is unclear which CT is contributing to the identified DNAm associations. We addressed this question with a stratification approach based on CT composition and our findings could have implications on informing future research design (e.g., what tissue type may be more sensitive to certain variable of interest), hypothesis formations for tissue or CT-specific effect, and replication of DNAm findings. Overall, our results showed that stratifying samples by BEC proportions for EWAS had differential effects across variables of interest, such that a larger number of significant DNAm associations were detected with both biological sex and cotinine concentration in stratified samples than full sample, but not with SES.

We showed that sample stratification allowed for detection of differential sex associations across CTs. However, the differential associations with sex across the three saliva subsamples were not as prominent as across matched blood and buccal samples in an independent cohort. Despite this, the DNAm sites consistently observed across BEC subsamples overlapped with all five CMRs that were previously reported to be associated with sex across multiple tissues, including buccal and blood tissues^27^. Hence, the concordance across stratified salivary samples parallels that of cross-tissue similarities.

We identified three significant DNAm associations with cotinine concentrations in our full sample, out of which two (cg05549970 and cg14588422 annotated to the *C4orf50* and *PARD3* genes, respectively) have been reported to be associated with first-hand smoking^28,29^. After stratification by BEC proportions, we identified significant DNAm associations with salivary cotinine concentrations only in the low BEC subsample. Despite a smaller sample size, low BEC subsample, which had a more homogeneous and immune-focused CT composition, showed a larger number of significant cotinine-DNAm associations than that identified in the full sample, reflecting potential CT-specific effects of cotinine on DNAm.

We did not find significant DNAm associations with SES at any site after stratification, yet the directionality and strength of effects at the significant sites identified in the full sample were strongly correlated in stratified samples. Therefore, a possible explanation for the absence of significant findings in CT-stratified samples is that with a smaller effect size than biological variable, the reduction of sample size may have decreased the power to detect the subtle effect. Furthermore, unlike cotinine, SES may not have a strong immune CT effect in saliva samples, or required further CT specificity than could be achieved by stratification. However, past studies have found differential associations across tissues with sociodemographic variables. For example one study comparing DNAm associations in buccal and blood samples indicated that blood DNAm had stronger associations than buccal DNAm^30^. In our saliva samples, the low BEC subsample, which has the highest immune cell proportion, had the smallest sample size and therefore, the least power to detect significant SES-associations if they were present. As such, it is unclear whether the lack of significant SES-associations in the low BEC subsample was due to the small sample size or the inherent difference between blood and saliva with high immune cell proportions given the differences in oral immune cells, which cannot be tested with our current sample but should be investigated in future studies.

### Differential EAA associations with variables of interests when considering BEC proportions in EAA calculations or stratifying samples by BEC proportions

Both PedBE, specifically trained in pediatric cheek swab samples, and Horvath Skin-blood clocks, trained in adult cells and samples including saliva, showed significant differences in EAA across stratified samples by BEC proportions. Specifically, children with lower BEC proportions had higher EAA. These EAA differences with the same directionality have been reported in previous studies on EAA in pediatric cheek swab samples^10^ and are now extended to pediatric saliva samples in our current study. This could be in part due to the computational approaches used to create epigenetic clocks that may select DNAm sites that are informative to age-related cell-type differences^16^. Importantly, these differences became non-significant in both clocks when EAA calculations accounted for CT proportions, suggesting the reduction of confounding effect and further supporting the existing recommendations to correct for estimated CT proportions when calculating EAA^10,16^. Overall, when we investigated EAA associations with our variables of interest, only PedBE, but not Skin-blood, EAAs produced significant findings. It is tempting to speculate that PedBE is more sensitive to signals in children than Skin-blood due to its training samples matching with our sample’s developmental stage.

Sex differences in PedBE EAAs were robust across methodological approaches—in both full and stratified samples as well as when EAAs were calculated accounting for CT proportions or not. Specifically, we found lower PedBE EAA in boys than girls, which is consistent with previous EAA findings in pediatric cohort^31^. It is important to note that the effect sizes of these sex differences became larger after stratification, potentially due to the reduced CT heterogeneity.

Extending our immune-specific cotinine-EWAS findings, we further showed that cotinine-EAA associations were also specific to stratified samples containing primarily immune cells. Together, these findings showed converging immune-specific effect across multi-modal DNAm analyses. The lower EAA associated with higher cotinine levels in the low BEC subsamples may seem counterintuitive given that higher biological aging is often found in adult smokers^32^. However, the interpretation of pediatric EAA is much less clear that adult EAA; more consistent links between higher EAA and aging phenotypes including disease and mortality has been established in adult literature^33^. Due to the relative scarcity of pediatric EAA studies, it is still unclear how slower pediatric EAA is linked to the spectrum of developmental outcomes. However, early psychosocial adversity is linked with lower gestational and pediatric EAA^34^. It is possible that any deviation, regardless of faster or slower EAA, can be indicative of non-normative development^35^.

When CT proportions were accounted for in EAA calculation, its associations with SES were amplified and became statistical significance; specifically, lower SES were associated with higher EAA. Previous literature has shown that early childhood SES were linked to higher EAA in adults^36,37^ and our finding demonstrated that such association may emerge early in life. Again, aligning with our EWAS on SES findings, no significant associations were found after stratification, yet similar trends were identified in both low and mid BEC subsamples. It is tempting to speculate that EAA associations with SES may be consistent across saliva samples regardless of CT compositions and hence stratification may not provide further insight to CT-specific SES associations.

### Limitations & Recommendations

The findings from the current study should be interpreted while acknowledging its limitations. First, our study did not include cell count data to compare the accuracy of CT predictions across reference panels. Our investigation focused on the impact of differential CT predictions on downstream analyses and how stratification based on these estimated CT proportions could illuminate interpretation. Given the heterogeneity in saliva samples across cohorts, it is crucial to first assess the degree of CT heterogeneity in the given saliva samples; with lower levels of CT heterogeneity, stratification of samples may not be as informative. We note that alternative approaches, such as CellDMC^38^ and eFORGE^38^, are also available to unpack the origin of DNAm signals in heterogeneous samples. However, given our saliva samples are primarily made up of BEC and immune cells and with clear subsamples based on primary CTs, we opted for a stratification approach to provide more interpretable results. Second, some of our samples had salivary cotinine concentrations that were too low to be reliably measured by the assay, which led to missingness in our data. This was not unexpected given the young age of our sample. Importantly, our main findings were supported by sensitivity analyses that used imputed cotinine concentration for samples with censored, indicating the robustness of our findings. Given that the main objective of the current study was to examine how different methods might affect DNAm analyses using cotinine concentrations as an example, we did not intend to draw conclusions from the significant DNAm sites identified in this study. Lastly, our study employed stringent QC to ensure high-quality DNAm samples for our investigation, which removed a substantial portion of samples. However, the QC process is cohort-dependent and can be influenced by many factors, including the extent to which child participants were able to follow instructions during sample collection and their oral health conditions, such as tooth decay, tooth loss, and cold sores^5,39^. Despite these challenges, saliva still presents a wide range of strengths in epidemiological research including its minimally-invasive sampling methods and opportunities for interdisciplinary research ranging from stress hormones to immune markers, from oral analytes to microbiome, in addition to genetics and epigenetic research^6^.

To inform future epigenetic research using pediatric saliva, we summarized the implications of our approaches with respect to the high CT heterogeneity in saliva in **Table 1**. In conclusion, CT references built on a population and tissue matched with the targeted samples produced more accurate estimations and the discrepancies in CT estimations had differential effects on DNAm associations with commonly investigated variables in the field. Further, stratification of samples by CT proportions illuminates the CT populations most sensitive to DNAm and EAA associations for suitable variables of interest. The current study contributes to future pediatric DNAm research by offering practical guidance to leverage this complex yet highly accessible biological sample in children.

**Table 1.** Implications for analytical approaches with pediatric saliva DNAm.

| Reference panel selection |  | Sample selection |  |
| --- | --- | --- | --- |
| Pediatric reference panel | Adult reference panel | Full sample | Stratifications by primary CT |
| • More appropriate CT estimation with pediatric reference<br>• More power to detect significant DNAm associations with social variable with pediatric reference | • Suitable for CT adjustments in EWAS with biological variables<br>• More comparable with past saliva DNAm analyses using the adult reference | Allow for identification of generalizable saliva biomarkers both in the context of EWAS and EAA | Enhance interpretability and replicability of salivary DNAm findings for variables of interest by identifying what primary CT has higher sensitivity |

*Note.* BEC=buccal epithelial cell; CT=cell type; DNAm=DNA methylation

## Supporting information

Supplementary Methods & Analyses

## Acknowledgements

We thank Kaitlin Smith, Hillary Piccerillo, Tatum Stauffer, and Andrew Huang for technical assistance with salivary biospecimen testing. We would like to express our gratitude to all of the families, participants, and teachers who participated in this research and to the Family Life Project (FLP) research assistants for their hard work and dedication to the FLP. This study is part of the Family Life Project. We thank the late Clancy Blair for his leadership of the Family Life Project (FLP), mentorship, and contributions to the early stages of this project. We thank Christopher Bartlett and his team for their technical support on the genotyping of the salivary samples.

