## Supplementary Methods & Analyses for "Not All Saliva Samples Are Equal: The Role of Cellular Heterogeneity in DNA methylation and Epigenetic Age Analyses with Biological and Psychosocial Factors"

**Supplementary material for Methods & Analyses**

**Participants**

There were no statistically significant differences between the FLP sample at 2 months, which was the time point with the most complete data (n=1292), and our current sample (n=529) in percentage of low-income families (75.2% in current sample vs. 77.6% in whole sample, p=.313), mother-reported child race as Black/African American (39.7% in current sample vs. 42.6% in whole sample, p=.330), and child biological sex (current sample=49.9% vs. whole sample=49.1% male, p=.644) based on chi-square tests. There was a small but statistically significant difference in age (current sample=7.26 vs. whole sample=7.29 years, p=0.03) between the whole and current samples.

**Collection of salivary samples**

Unstimulated whole saliva samples were collected from children using the passive drool method (Granger et al., 2012) during the 90 month visit to participants’ homes. At the time of collection, samples were frozen at -20^◦^C and then transferred to the Institute for Interdisciplinary Salivary Bioscience Research at the University of California, Irvine, CA, USA for archiving at -80^◦^C. At the time of use, saliva samples were thawed and centrifuged (5,000 g; 10 min; 4^◦^C) to remove insoluble material and cellular debris. The resulting cell pellet was stored at -80°C until DNA extraction.

**DNA extractions**

Pellet fractions from the centrifuged saliva samples were resuspended in 200 µl of PBS. DNA was then isolated using the QIAamp DNA Mini Kits (QIAGEN, Cat #1304, Hilden, Germany) following the manufacturer’s instructions, with the exception that twice as much Proteinase K was used and samples were incubated with Proteinase K for 2 hours. DNA quality and quantity was determined using a NanoDrop Spectrophotometer (ThermoFisher, Waltham, MA, USA).

**DNAm preprocessing**

Briefly, 750 ng of genomic DNA was bisulfite-converted using EZ DNA Methylation Kit (Zymo Research, CA, USA). Subsequently, 160 ng of bisulfite-converted DNA was applied to the EPIC array, following manufacturer’s protocols (Illumina, CA, USA). Processed EPIC arrays were scanned with an Illumina iScan (Illumina, CA, USA). QC of the EPIC array DNAm data was performed in R (4.2.2) using a combination of the minfi package (v1.44.0) (Fortin et al., 2017), which plots the log median intensity of a sample in both the methylated and unmethylated channels, and ewastools (v1.7.2) (Murat et al., 2020), which provides seventeen QC metrics as described by Illumina in their BeadArray Controls Reporter Software Guide document packages. A total of 200 samples were removed after all QC steps (see **Supplementary Figure S1a** for details). One possible reason for this high number of removed samples is bacterial contamination. Bacterial DNA assay was run on the failed samples with remaining saliva available and showed that lower cycle threshold (Ct) values (i.e., faster amplification) (mean = 10.95) than samples that passed QC (mean = 11.71), *t*(24) = 3.08, p = 0.005 (**Supplementary Figure S1a**).

Our additional samples with age mismatch (>10 years difference) based on the epigenetic age calculated with the Pediatric Buccal (PedBE) (McEwen et al., 2020) and Pan-tissue Horvath clocks (Horvath, 2013) in the methylclock R package and reported chronological age were removed. These age mismatches were likely due to mislabelling of saliva samples from the child and caregiver (**Supplementary Table S1**). After sample removal, both PedBE and Horvath Skin-blood epigenetic ages were statistically significantly and positively correlated with chronological age (*p*s = 9.599e-08 and 7.404e-09), with PedBE showing slightly stronger correlation and smaller mean absolute error (MAE) (r = 0.25, MAE = 0.95 years) than Skin-blood (r = 0.23, MAE = 2.06 years). Given the small age range in our samples, MAE may be more informative than the correlation coefficient.

A total of 529 high-quality samples were included for downstream analyses (**Supplementary figure S1**). The removed samples did not differ from the final sample based on demographic variables, including child sex, low-income status, and parent-report child ethnicity (*p*s = 0.601 - 0.962).

**Supplementary Table S1**. Details of the four samples with age mismatch

| **Child reported Age** | **Child sex** | **PedBE epigenetic age** | **Horvath epigenetic age** | **Caregiver reported age** | **Caregiver sex** |
| --- | --- | --- | --- | --- | --- |
| 7.28 | F | 26.89 | 33.62 | 28.98 | F |
| 7.14 | F | 23.96 | 24.73 | 26.45 | F |
| 7.37 | F | 29.22 | 21.51 | 28.62 | F |
| 7.22 | F | 30.20 | 34.23 | 39.66 | F |

To account for type I and type II probe differences on the EPIC array, DNAm data was normalized using functional normalization by *minfi::preprocessFunnorm*. Probe filtering was conducted on the normalized DNAm data (see **Supplementary** **Figure S1b**). In total, 41,787 cross-reactive probes were removed (Pidsley et al., 2016). Next, polymorphic probes containing a single nucleotide polymorphism (SNP) at the target DNAm site (n probes = 11,270) or at the base extension sites (48 and 49 base pairs for Infinium Type I and II probes, respectively) (n = 259) with minor allele frequency (MAF) >5% (Pidsley et al., 2016), based on the 1000 Genomes project (Siva, 2008), including genotypes from 26 different populations, were also removed. Then, DNAm sites which overlapped with probes located on the sex chromosomes, and probes with a detection p-value > 0.01, as well as those with NAs in more than 2% of samples using *watermelon::pfilter* were also removed*.* A total of 786,069 probes passed the QC. We performed *sva::ComBat* to account for remaining technical effects on plate and chip (E. M. Price & Robinson, 2018).

**Whole genome genotyping & genetic principal components**

QC was conducted as described in Simmons et al. (2011), including individual/SNP genotype characteristics, inferring biological relations, and ancestry characteristics. Briefly, SNPs with MAF < 1%, SNP-level missingness > 1% or a Hardy-Weinberg equilibrium test with *p* <1x10^-6^ were filtered. Individual samples with excess/reduced heterozygosity, individual-level missingness > 1%, or incorrect sex determination were filtered.

We analyzed population ancestry using principal components analysis (PCA) calculated with the smartpca function from the EIGENSOFT package that computes PC scores from SNPs (A. L. Price et al., 2008). We analyzed all samples that passed QC along with all 1000 Genomes samples (Siva, 2008) to provide clear reference populations for the major continental groupings. To adjust for effects of genetic admixture, we included the first three PCs to in all analyses.

**Salivary cotinine measure**

This assay has a test volume of 20 µL, range of standards from 0.8 to 200 ng/L, and lower limit of detection (LLD) of 0.15 ng/L. Cotinine high and low controls were run on every assay plate. The intra-assay and inter-assay coefficients of variation (CVs) were 7.1% and 9.1%, respectively. A concentration below 10 ng/ml is indicative of potential passive environmental tobacco exposure (Rao et al., 2023), which is in line with our observation in this cohort (mean = 2.53 ng/ml).

**Characterizing and comparing CT proportion estimates across adult and pediatric saliva reference panels**

For the adult reference, three major subsets (epithelial cells, fibroblasts, and immune cells) were first deconvoluted with the *EpiDISH::*centEpiFibIC.m reference matrix. Subsequently, we used *EpiDISH::hepidish* to estimate seven CT proportions based on epigenomic deconvolution (B-cells, CD4+ T-cells, CD8+ T-cells, NK cells, monocytes, neutrophils, and eosinophils) with the *EpiDISH::*centBloodSub.m reference matrix. As for the pediatric reference, the saliva samples were first deconvoluted into the BEC versus immune cell proportions, also followed by deconvolution of the immune cells with the *EpiDISH::*centBloodSub.m reference matrix.

We assessed the median and interquartile range (IQR), the variance and distributions of estimated BEC proportions, and the appropriateness of the reference panels using the CEll TYpe deconvolution GOodness (CETYGO) score (*CETYGO*::projectCellTypeWithError), a metric to assess the accuracy of cellular deconvolution when actual cell count is not available. The CETYGO score was defined as the root mean squared error (RMSE) between the measured DNAm profile in the sample and the expected profile from the reference panels. Therefore, a perfect estimate would have a CETYGO score of 0 (i.e., lower scores are more accurate), while higher values reflect less accurate estimations of CT compositions. A CETYGO score<0.1 indicates an appropriate reference, while a score>0.1 suggests the reference panel may not be relevant for the tissue being profiled

To reduce the dimensionality of our CT proportion variables, we performed isometric log-ratio transformation on the estimated CT proportion, followed by robust principal component analysis (PCA) with robCompositions::pcaCoDa. Given that our findings demonstrated appropriateness of CT proportions estimated by the child reference, we stratified our samplesmusing the child-based estimated CT proportions. We first characterized the three subsamples stratified by estimated BEC proportion before comparing the DNAm associations across these subsamples (**Supplementary** Table 1). We did not find any significant differences across demographic variables, including child sex, age, parent-reported race, SES, and cotinine concentration based on one-way ANOVA, *p*s=0.38-0.97.

**Epigenome-wide association study (EWAS): Delta beta calculation**

In our analyses for biological sex (a binary variable), Δβ was the regression coefficient of the variable from the robust linear regression described above. For cotinine concentration and SES (continuous variables), Δβ was calculated by extracting the regression coefficient of the variable of interest from the robust linear regression models described above and then multiplying the coefficient with the range of cotinine or SES values between the 5th and 95th percentiles to reduce the effects of outliers. The resulting Δβ value represents a change in DNAm β value at each site associated with cotinine concentration or SES while adjusting for other covariates.

**Sensitivity analyses for cotinine measures**

To address the potential effect of the missing salivary cotinine data on our results, sensitivity analyses were conducted to address the censored data below the assay’s LLD (0.15ng/L) by imputing these data with half the LLD of the assay (i.e., 0.075ng/L; total n for these analyses=523 with mean=1.93ng/L, range=0.075-47.08ng/L). Furthermore, we categorized the cotinine concentration into a three-level variable of low exposure (n=235; ≤0.45 ng/mL), moderate exposure (n=279; 0.46–12 ng/mL), and high exposure (n=9; ≥12 ng/mL) as described in a previous study^25^. We also conducted sensitivity analyses on low vs. moderate exposure to test the robustness of our findings when the cotinine measure was operationalized differently.

**Validation of cross-tissue comparison for sex-DNAm associations**

We leveraged the 50 matched buccal and peripheral blood mononuclear cells (PBMC) samples from the GECKO cohort, a Canadian cohort of children aged 7-13 years old, previously described elsewhere (43.1% female, 72.5% White as reported by parents)^26,27^. EWAS on biological sex was conducted separately in the buccal and PBMC samples while adjusting for similar covariates as the primary analyses. Significant associations were determined by the same thresholds across tissues (statistical: FDR<.05 and technical: |Δβ|>.05). The number of overlapping significant DNAm sites across the GECKO matched tissue samples was compared with those across the FLP stratified subsets.

**Permutation**

With the new subsets in each permutation, we ran the same EWAS as described above (i.e., 100 permutations for each of the three EWAS). The resulting value is the permutation p-value that represents the likelihood that the unique number of associations of each group as well as the number of overlapping associations across all groups were observed due to chance. When the BEC group with a larger sample size showed a higher number of significant associations, we also ran 100 permutations to randomly draw a sample matching with the size of the smallest group.

To assess whether the difference in the amount of statistically significant associations were driven by differences in sample sizes (given that the high BEC subsamples [n=195] was higher than the low BEC subsamples [n=123]), permutations were conducted. The results suggested no significant reduction in the number of DNAm associations found in the high BEC group when 123 samples (same sample size as the smallest BEC group size) were randomly drawn from the 195 samples (median n associations=1,009; *p*=0.14).

We tested whether it is likely to find a percentage of overlapping sex associations across all stratified samples of 11.9% by chance. We performed 100 randomized permutations, which showed that a percentage of overlapping associations across the three groups as low as or lower than 11.9% were not due to random grouping of samples (*p*=0.03; median=26.00%; range=5.22-36.45%). Similarly, the unique number of associations (3,007 sites) in the high BEC was unlikely to be observed by random grouping (*p*=0.03).

One hundred randomized permutations indicated that the number of unique statistically significant associations with cotinine concentration in the low BEC subsamples was not significantly higher than by chance (p=0.09).
