## Supplementary Figures for "Not All Saliva Samples Are Equal: The Role of Cellular Heterogeneity in DNA methylation and Epigenetic Age Analyses with Biological and Psychosocial Factors"

**Supplementary materials**

**Supplementary Table S1.** Full sample (n=529) and stratified samples characteristics

**Supplementary Figure S1.** DNA methylation preprocessing

**Supplementary Figure S2.** Cell type predictions using an adult (upper panel) and child (lower panel) reference dataset.

**Supplementary Figure S3.** Similar results for EWAS across child and adult reference panels on biological sex and cotinine concentration but different on socioeconomic status.

**Supplementary Figure S4.** Sensitivity analyses of cotinine variables across child and adult reference panels.

**Supplementary Figure S5.** Significant DNAm associations with biological sex across all three BEC subsets, salivary cotinine concentrations in low BEC groups, and no significant association with SES.

**Supplementary Figure S6**. Significant associations with sex across buccal epithelial cells and peripheral blood monocytes samples in the GECKO cohort.

**Supplementary Figure S7.** Sensitivity analyses of stratified samples DNA methylation associations with cotinine variables.

**Supplementary figure S8** Strong correlations between delta betas of significant SES-associated DNAm sites in full sample and stratified samples.

**Supplementary Figure S9** Horvath Skin-blood EAA associations with biological sex, cotinine concentration, and SES across BEC stratified subsets.

**Supplementary Table S1**. Full sample (n=529) and stratified samples characteristics

| **Variables** | **n** | **Mean** | **SD** |
| --- | --- | --- | --- |
| Age (years) | 529 | 7.26 | 0.26 |
| Low BEC | 123 | 7.22 | 0.30 |
| Mid BEC | 211 | 7.24 | 0.24 |
| High BEC | 195 | 7.27 | 0.25 |
| Biological sex (male) | 529 | 49.9% | – |
| Low BEC | 123 | 52.8% | – |
| Mid BEC | 211 | 49.3% | – |
| High BEC | 195 | 47.7% | – |
| Family income-to-needs ratio | 529 | 1.91 | 1.61 |
| Low BEC | 123 | 1.80 | 1.64 |
| Mid BEC | 211 | 2.03 | 1.74 |
| High BEC | 195 | 1.83 | 1.43 |
| Salivary cotinine (ng/mL) | 394^a^ | 2.53 | 3.74 |
| Low BEC | 101 | 3.01 | 5.41 |
| Mid BEC | 159 | 2.14 | 2.54 |
| High BEC | 134 | 2.63 | 3.36 |

*Note*. SD=standard deviation; BEC= buccal epithelial cell; SES=socioeconomic status; ^a^the smaller sample size since concentrations below the lower limit of detection (n=135) were treated as missing data in the main analyses.

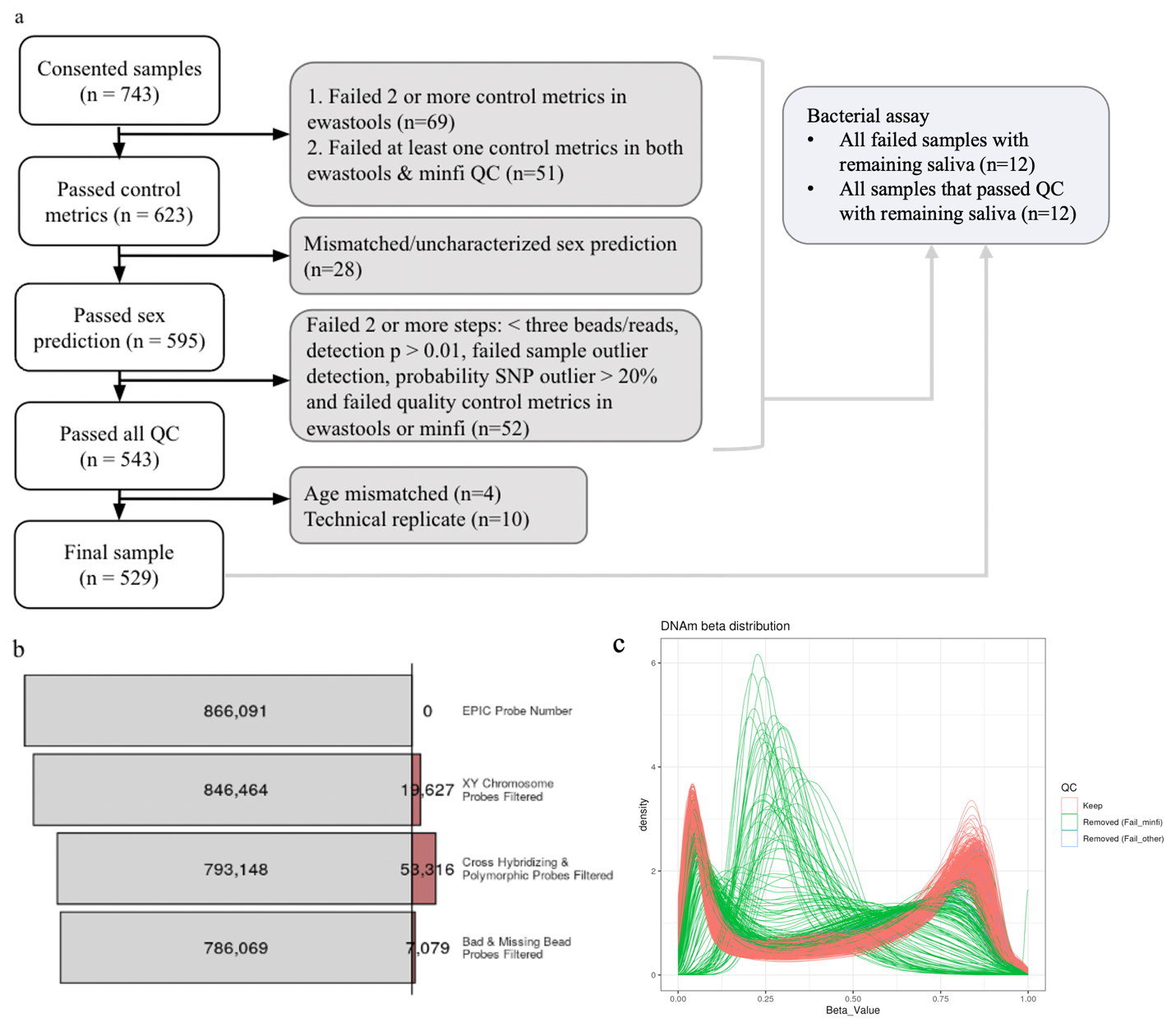

**Supplementary Figure S1 DNA methylation preprocessing.** Quality control (QC) was performed on the DNA methylation data. a) QC was performed on a total of 743 samples from whom consent was received, and the number of samples failed at each step. A final high-quality sample of n=529 individuals that passed all QC steps were included in the analyses in the current study, b) The number of probes filtered at each filtering step. A total of 786,069 probes remained after probe filtering, and c) Beta distribution of DNA methylation sites from samples that have passed the QC (red lines) and that have failed QC (green and blue lines)

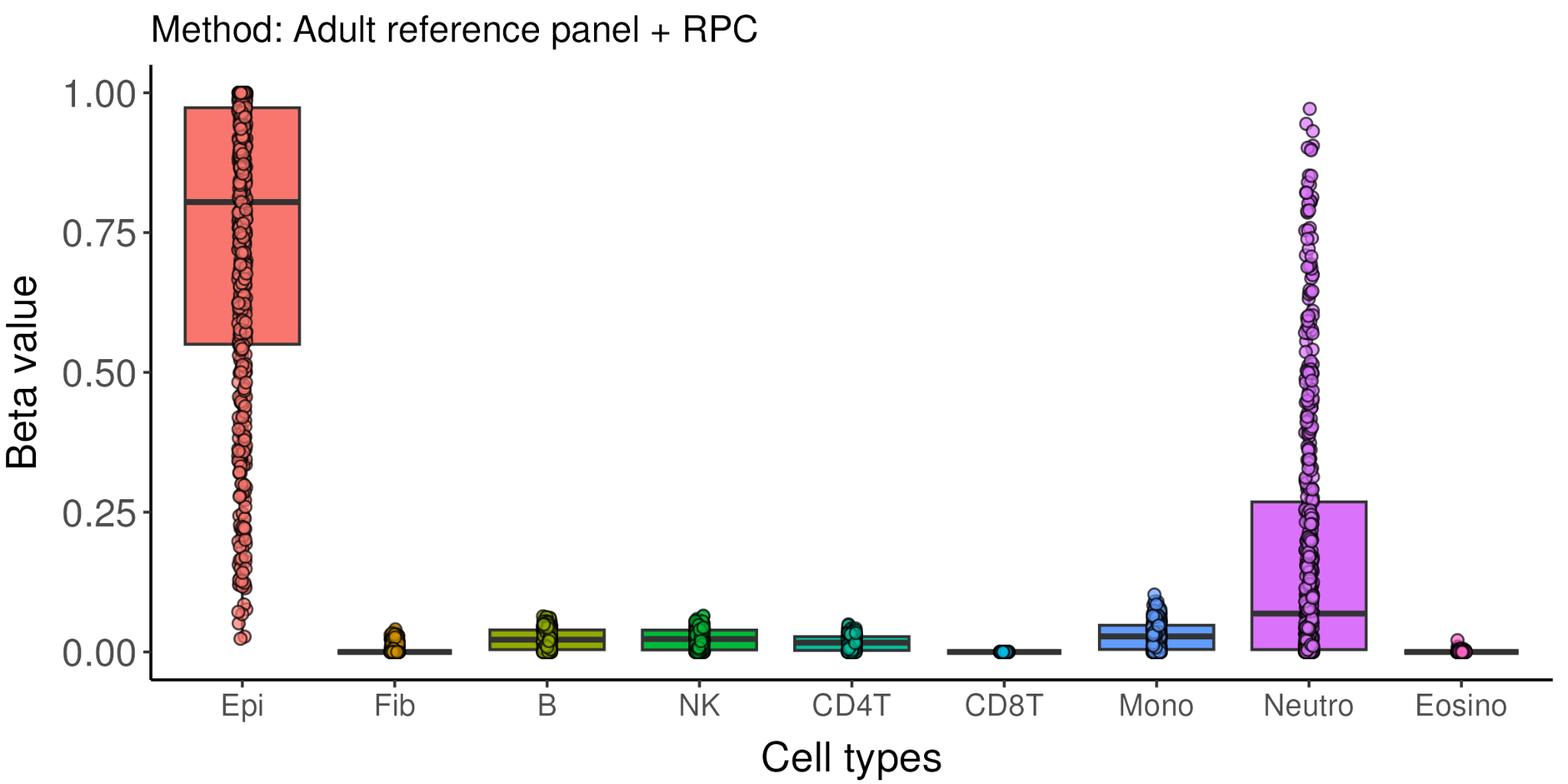

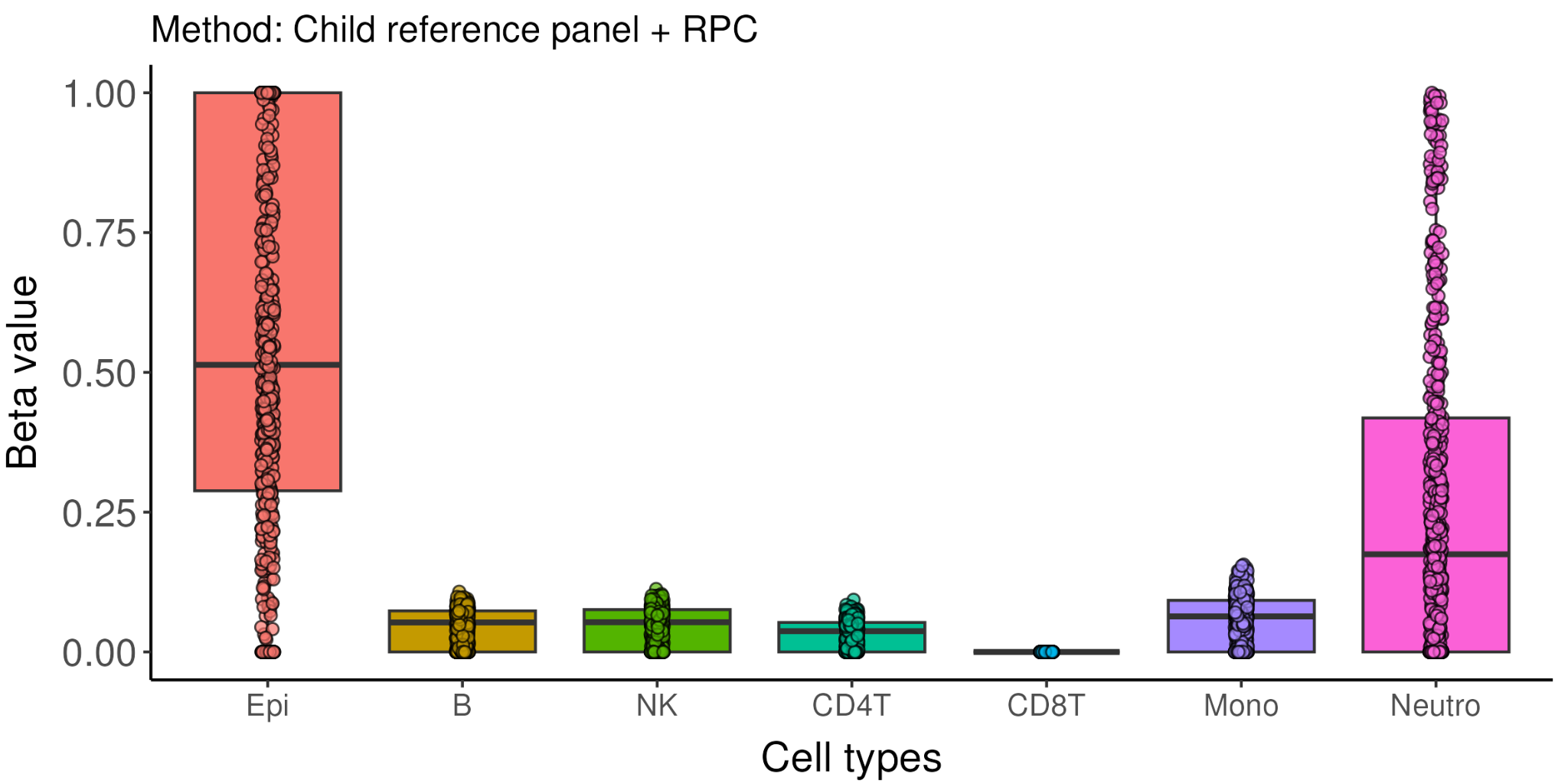

**Supplementary Figure S2. Cell type predictions using an adult (upper panel) and child (lower panel) reference dataset.**

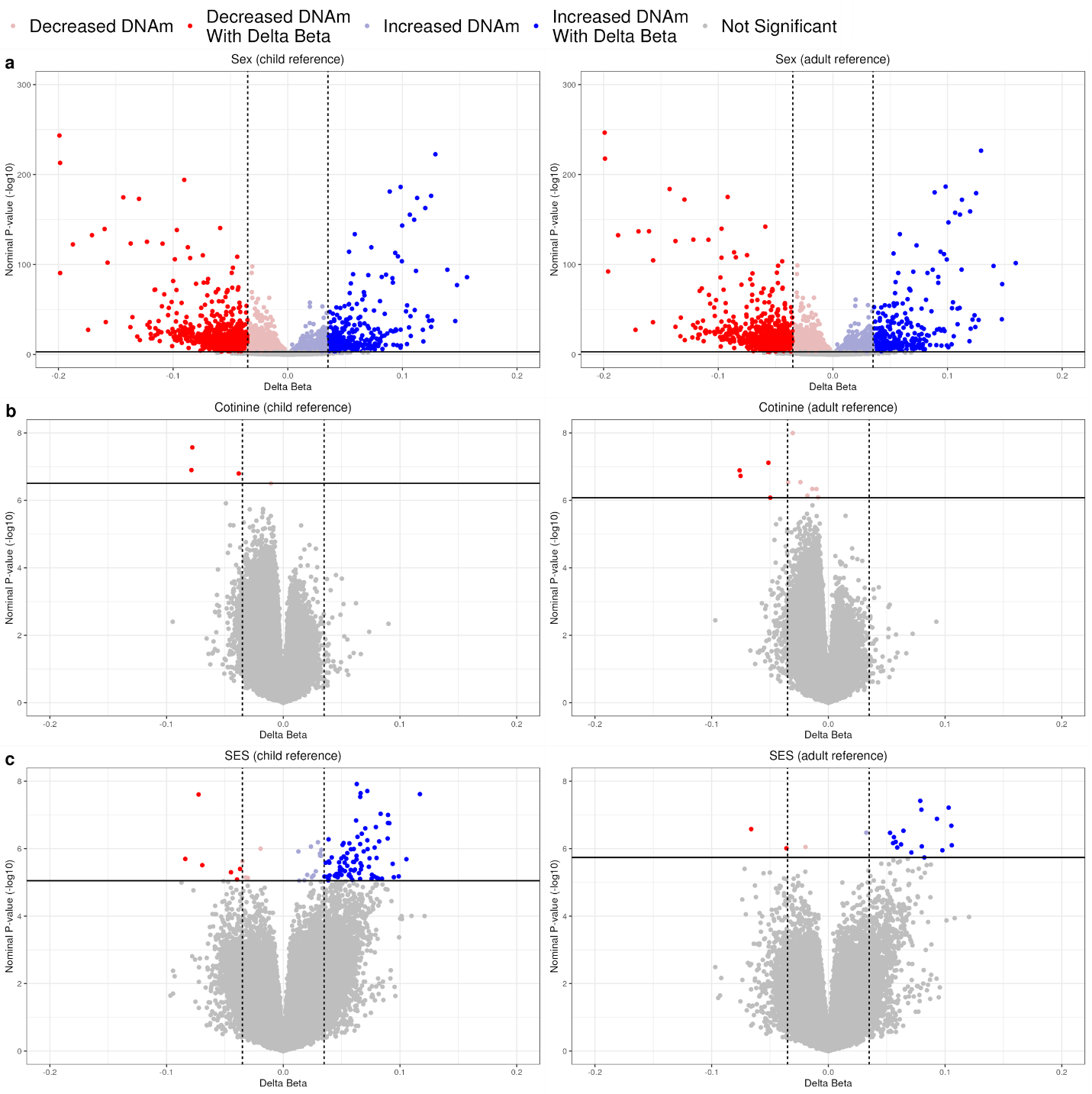

**Supplementary Figure S3. Similar results for EWAS across child and adult reference panels on biological sex and cotinine concentration but different on socioeconomic status.** Volcano plots of EWAS on a) biological sex, b) cotinine concentration, and c) socioeconomic status. The colored dots represented significant DNA methylation associations with biological sex, passing the statistical threshold of FDR < .05 and |Δβ| > .035. Red dots represent DNAm sites with a negative association, whereas blue dots represent DNAm sites with a positive association.

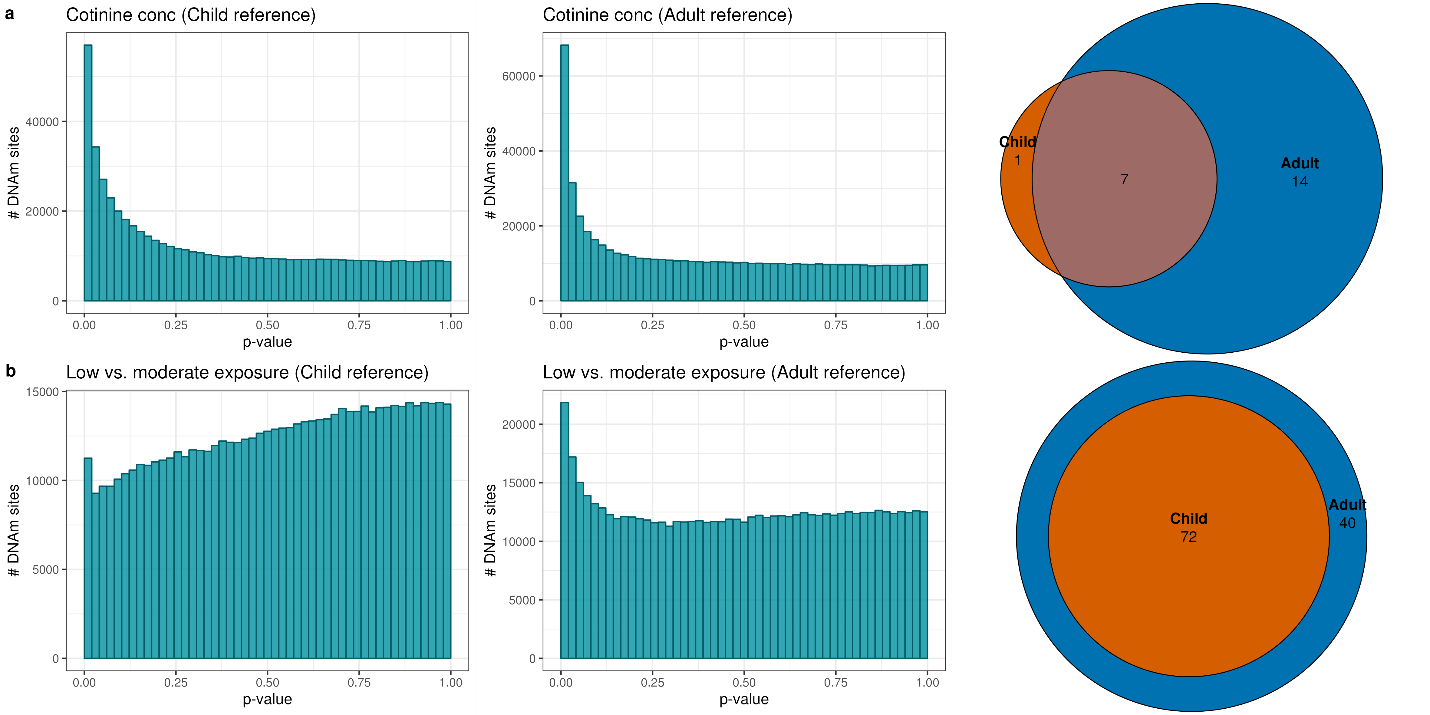
**Supplementary Figure S4. Sensitivity analyses of cotinine variables across child and adult reference panels.** Conc = concentration. Each row showed the p-value histograms and Venn diagrams representing the unique and overlapping sites across the child and adult reference panels of each cotinine variable (a: cotinine concentration with imputed data, and b: categorical variable of smoke-exposure).

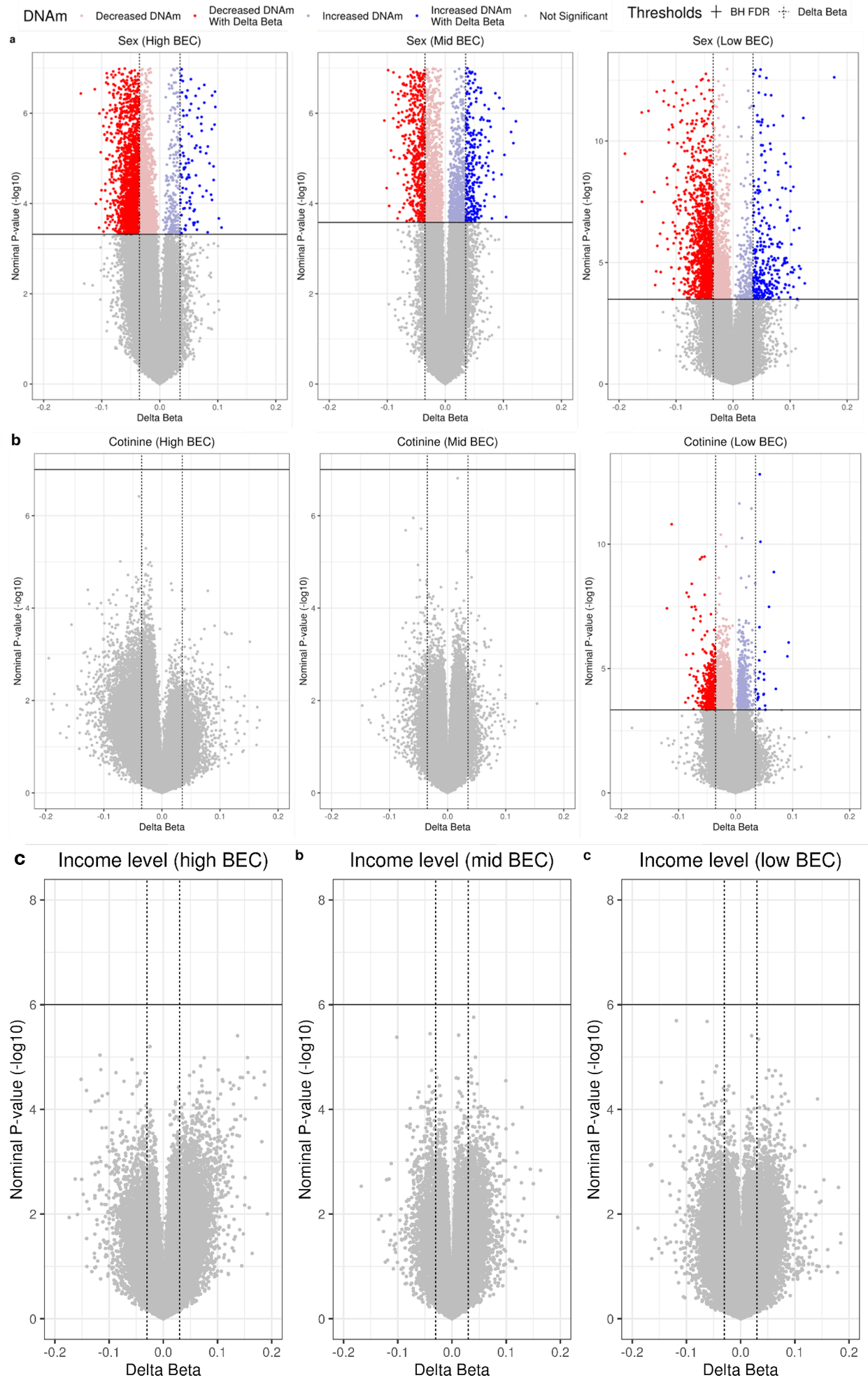

**Supplementary Figure S5 Significant DNAm associations with biological sex across all three BEC subsets, salivary cotinine concentrations in low BEC groups, and no significant association with SES.** Volcano plots displaying DNAm association with a) biological sex and b) salivary cotinine concentrations in the low, middle, and high BEC subsamples. The colored dots represented significant DNAm associations with biological sex, passing the statistical threshold of FDR<.05 and |Δβ|>.035. Red dots represent DNAm sites with a negative association, whereas blue dots represent DNAm sites with a positive association.

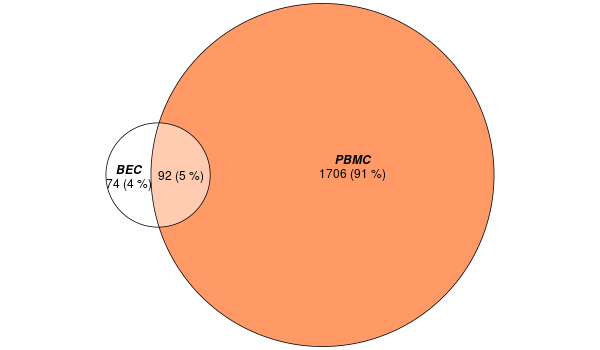

**Supplementary Figure S6 Significant associations with sex across buccal epithelial cells and peripheral blood monocytes samples in the GECKO cohort.** EWAS was conducted for sex in matched buccal epithelial cells (BEC) and peripheral blood monocytes (PBMC) samples in the GECKO cohort while adjusting for estimated cell type proportions. The Venn diagram depicts the overlapping and unique associations (number and percentage of total) across the two tissues. White circle (left) contains numbers for BEC, and the orange circle (right) contains numbers for PBMC.

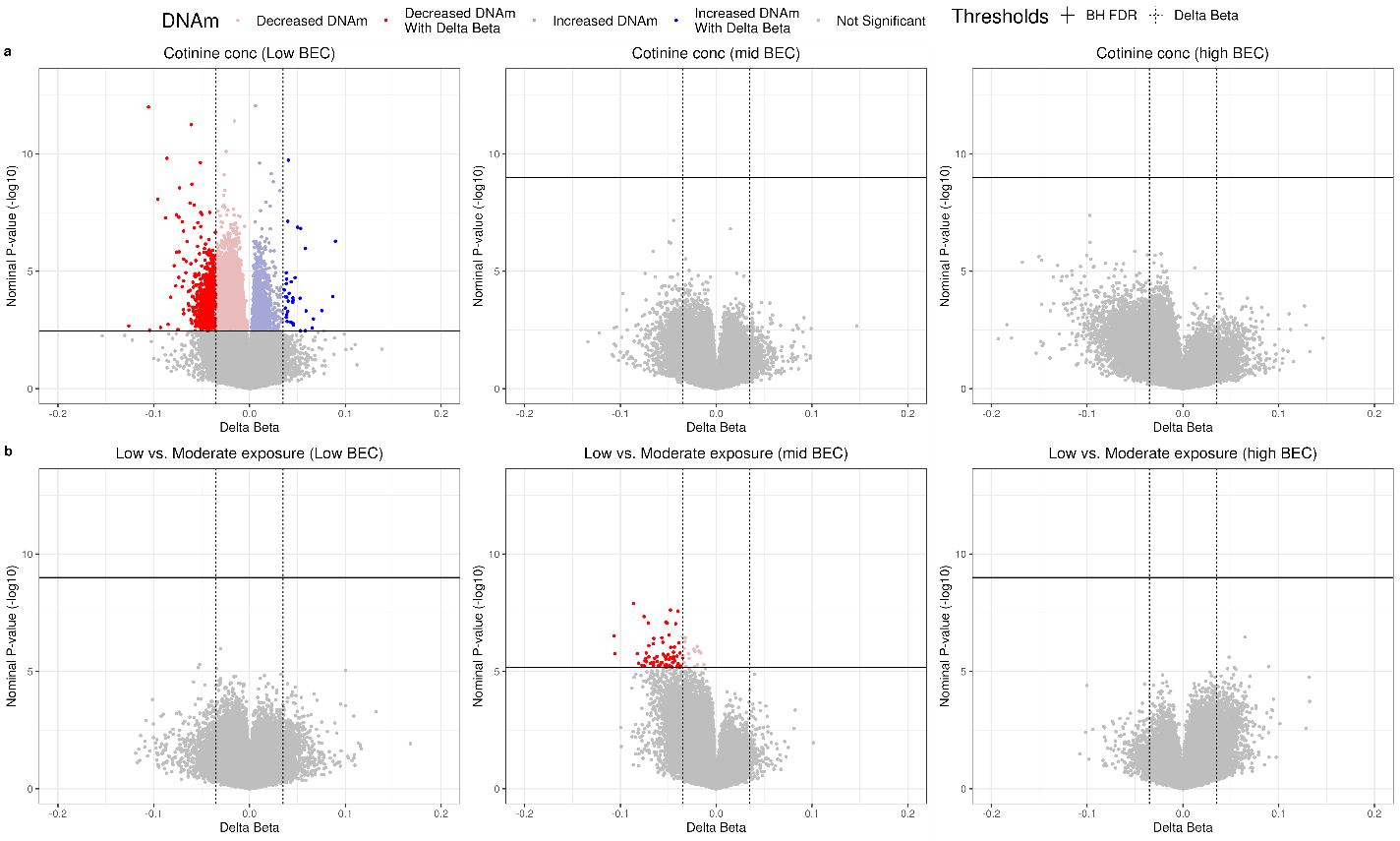

**Supplementary Figure S7. Sensitivity analyses of stratified samples DNA methylation associations with cotinine variables.** Conc = concentration. Volcano plots displaying DNAm associations with a) cotinine concentrations with imputed data and b) low vs. moderate exposure in the low, middle, and high BEC subsamples. The mean level of imputed cotinine concentration in the low, middle, and high BEC subsamples were 2.52 ng/mL, 1.65 ng/mL, and 1.86 ng/mL, respectively. The sample distribution of low and moderate smoke-exposure in the three BEC subsamples were as follow: n = 51 and 67 in low BEC group, 88 and 120 in middle BEC group, and 96 and 92 in high BEC group. Nine samples in the high smoke-exposure group were excluded from the sensitivity analyses. The colored dots represented significant DNA methylation associations with cotinine concentration, passing the statistical threshold of FDR < .05 and |Δβ| > .035. Red dots represent DNAm sites with a negative association, whereas blue dots represent DNAm sites with a positive association. The significant associations in the low BEC group were not found in the sensitivity analysis with the low vs. moderate exposure groups. It is worth noting that CT proportion PCs included in the EWAS models were significantly different across smoke exposure groups (t-test results, *p*s=.038 and .024) only in the low BEC group but not in other BEC groups. However, no significant correlations were found between CT proportion PCs and cotinine concentration variables (*r*s=-0.18–-0.13, *p*s=0.10-0.26).

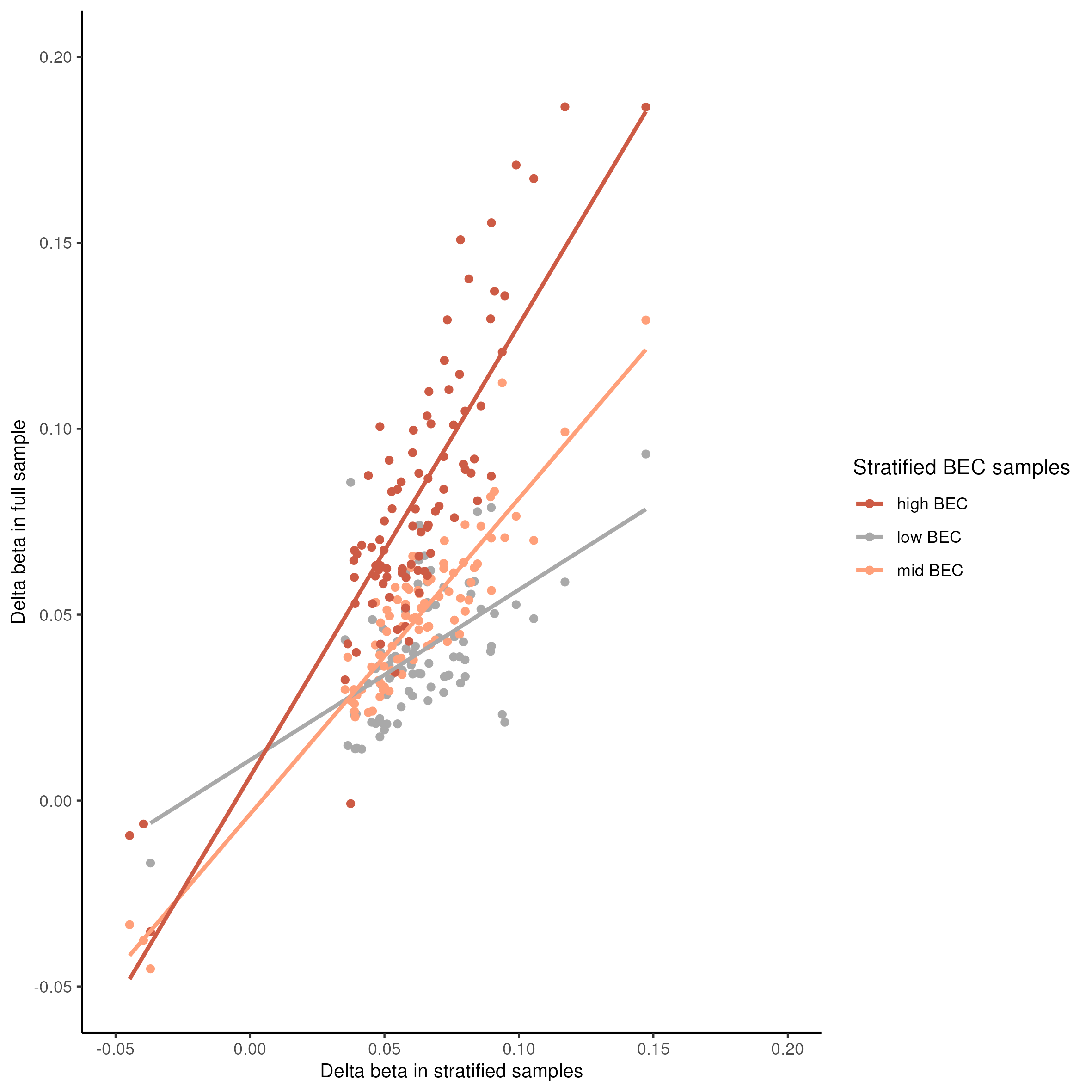

**Supplementary figure S8 Strong correlations between delta betas of significant SES-associated DNAm sites in full sample and stratified samples.** Scatterplot with regression line showing strong correlations between delta betas of significant DNAm sites in full sample and stratified samples. Brown dots and line represent high BEC subset, orange dots and line represent mid BEC, and grey dot and line represent low BEC. High BEC subset showed the strongest correlation with full sample.

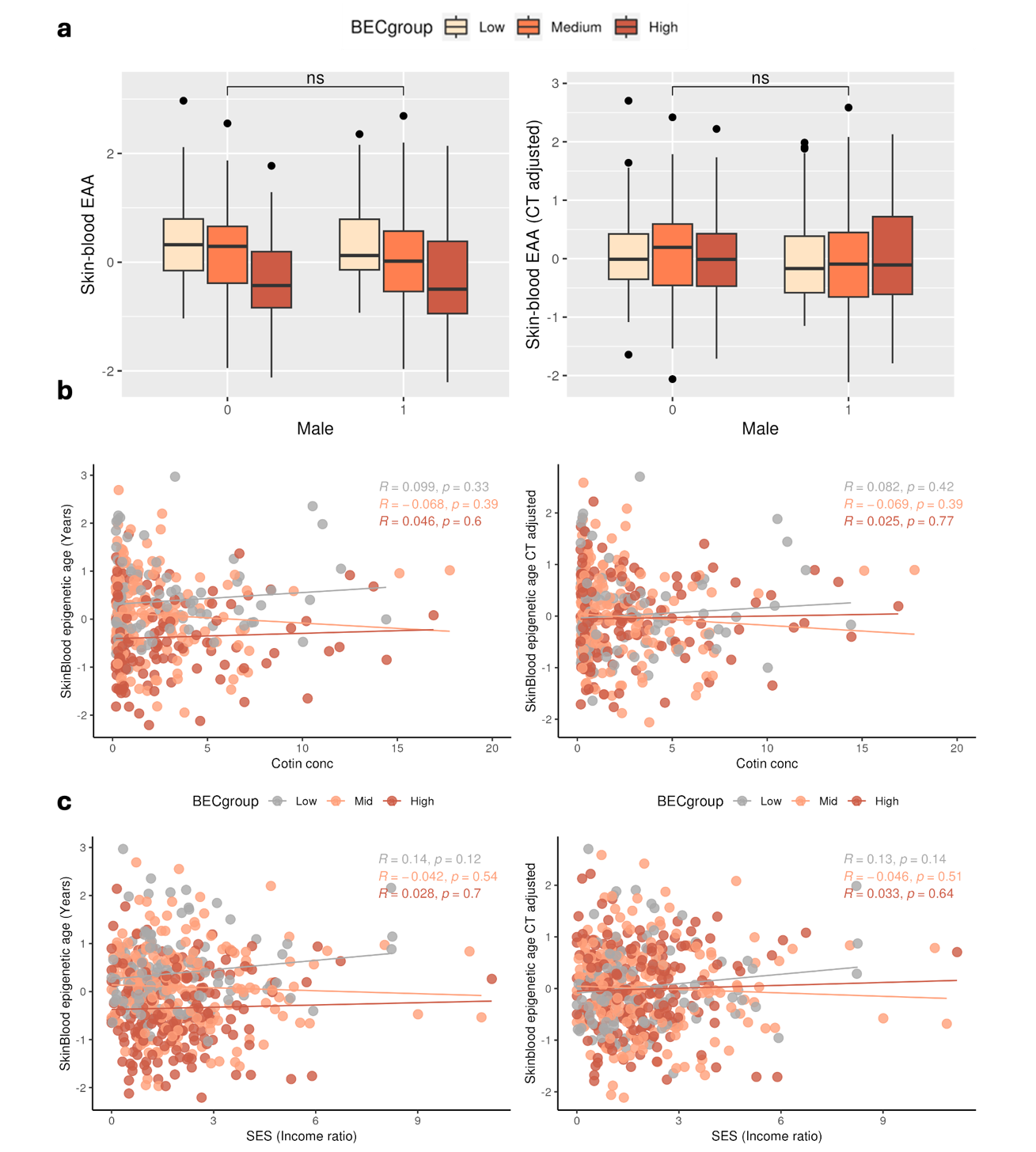

**Supplementary Figure S9 Horvath Skin-blood EAA associations with biological sex, cotinine concentration, and SES across BEC stratified subsets.** Boxplots in panel *a* showed no sex differences in Skin-blood EAAs (both with and without CT adjustment). Scatterplots in panel *b* and *c* showed that there were no associations between Skin-blood EAA (both with and without CT adjustment) with cotinine concentrations and SES.
